# Mitochondrial dysfunction and mitophagy blockade contribute to renal osteodystrophy in chronic kidney disease-mineral bone disorder

**DOI:** 10.1101/2023.12.26.573355

**Authors:** Shun-Neng Hsu, Louise A Stephen, Kanchan Phadwal, Scott Dillon, Roderick Carter, Nicholas M Morton, Ineke Luijten, Katie Emelianova, Anish K Amin, Vicky E Macrae, Tom C. Freeman, Katherine A Staines, Colin Farquharson

## Abstract

Chronic kidney disease–mineral and bone disorder (CKD-MBD) presents with extra-skeletal calcification and renal osteodystrophy (ROD). The origins of ROD likely lie with elevated uremic toxins and/or an altered hormonal profile but the cellular events responsible remain unclear. Here, we report that stalled mitophagy contributes to mitochondrial dysfunction in bones of a CKD-MBD mouse model, and also human CKD-MBD patients. RNA-seq analysis exposed an altered expression of genes associated with mitophagy and mitochondrial function in tibia of CKD-MBD mice. The accumulation of damaged osteocyte mitochondria and the expression of mitophagy regulators, p62/SQSTM1, ATG7 and LC3 was inconsistent with functional mitophagy, and in *mito*-QC reporter mice with CKD-MBD, there was a 2.3-fold increase in osteocyte mitolysosomes. Altered expression of mitophagy regulators in human CKD-MBD bones was also observed. To determine if uremic toxins were possibly responsible for these observations, indoxyl sulfate treatment of osteoblasts revealed mitochondria with distorted morphology and whose membrane potential and oxidative phosphorylation were decreased, and oxygen-free radical production increased. The altered p62**/**SQSTM1 and LC3-II expression was consistent with impaired mitophagy machinery and the effects of indoxyl sulfate were reversible by rapamycin. In conclusion, mitolysosome accumulation from impaired clearance of damaged mitochondria may contribute to the skeletal complications, characteristic of ROD. Targeting mitochondria and the mitophagy process may therefore offer novel routes for intervention to preserve bone health in patients with ROD. Such approaches would be timely as our current armamentarium of anti-fracture medications has not been developed for, or adequately studied in, patients with severe CKD-MBD.

**Graphical Abstract:** 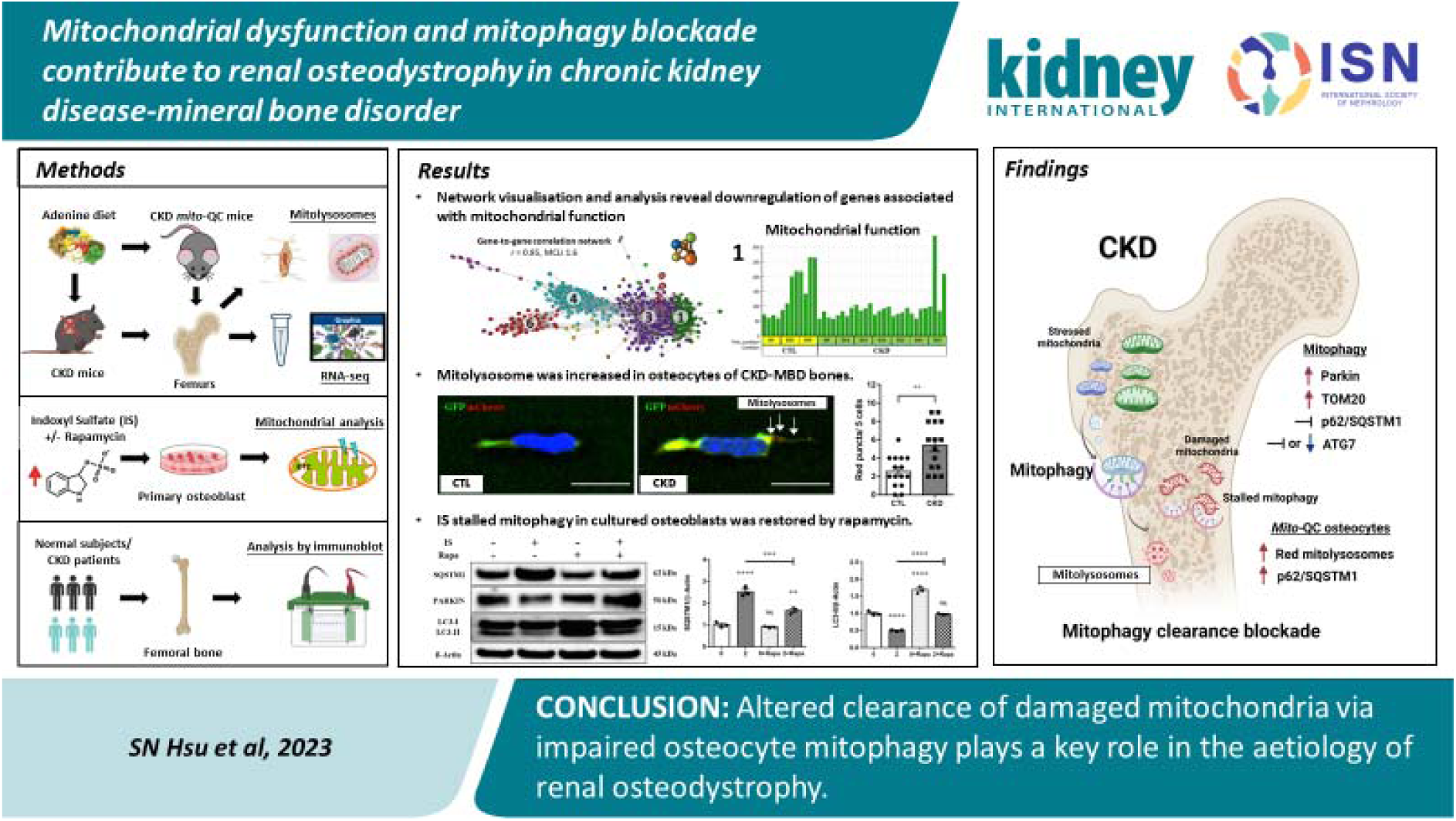

**TRANSLATIONAL STATEMENT:** Renal osteodystrophy (ROD) remains the major skeletal complication of chronic kidney disease-mineral and bone disorder (CKD-MBD). As a disease characterised by biochemical and hormone abnormalities, ROD is exacerbated by osteocyte mitochondrial dysfunction. Advances in our understanding of the mitophagy pathway are vital to improving the clinical management of ROD. The dysregulation of mitophagy in murine and human CKD-MBD bone provided evidence of delayed clearance of damaged mitochondria, which was also observed in uremic toxin-treated-osteoblasts but reversible upon rapamycin treatment. This study reveals the therapeutic potential of managing ROD by restoring defective mitophagy in osteocytes.

## INTRODUCTION

Chronic kidney disease-mineral and bone disorder (CKD-MBD) is a systemic disorder characterised by structural changes and the gradual loss of kidney function.^1^ Diagnosis is strongly correlated with the increasing prevalence of diabetes and hypertension, as well as an ageing population, and it is therefore a growing global health concern.^2^ Progressive loss of kidney function leads to secondary hyperparathyroidism (SHPT), hyperphosphatemia, and increased fibroblastic growth factor-23 (FGF23) levels. Through direct and indirect effects on bone-forming osteoblasts and bone-resorbing osteoclasts, this altered systemic milieu deregulates bone remodelling, mineral metabolism, and matrix mineralisation, leading to compromised bone formation and ectopic calcification.^3, 4^ The severe skeletal complications of CKD-MBD are numerous and, based on bone morphology, often classified into a wide spectrum of bone disorders generically referred to as renal osteodystrophy (ROD).^5^ Therapy often focuses on normalising defective bone remodelling, but despite this, ROD often results in bone fractures, negatively impacting life quality and the survival of renal patients.^6, 7^

While most CKD-MBD studies have focused on the perturbation of mineral-regulating hormones, including parathyroid hormone (PTH), FGF23, and vitamin D,^8^ the complex bone-renal signalling pathways and cellular events they control during the development of ROD are unclear. Nevertheless, it is recognised that SHPT, particularly common in late-stage CKD-MBD, is associated with a net loss of bone mass despite an accelerating bone turnover.^9^ This is a consequence of increased bone resorption by the osteoclast through the up-regulation of receptor activator of nuclear factor-κB ligand (RANKL) and the suppression of the RANKL decoy receptor, osteoprotegerin (OPG).^10, 11^ Retention of protein-bound uremic toxins such as indoxyl sulfate (IS) and p-cresyl sulfate in the circulation and organs of CKD patients can also affect bone quality.^12, 13^ Uremic toxins induce oxygen free radical production by mitochondria, and the resultant oxidative stress disrupts bone quantity and quality due to the unfavourable effects on osteoblasts and osteoclasts and the mineralisation process.^12, 14^ Uremic toxins also reduce the number of osteoblast PTH receptors, and the resultant skeletal resistance to PTH is likely to explain the aberrant bone turnover rate during the early stages of CKD-MBD.^15^

Our inadequate understanding of the cellular mechanisms implicated in the aetiology of renal bone disease, drove us to complete a series of studies aimed at providing insight into the mechanism(s) responsible. The identification of dysregulated mitophagy by uremic toxins and the accumulation of damaged mitochondria in osteocytes may help identify novel therapeutic options to manage skeletal complications in renal patients.

## METHODS

See the supplementary methods for detailed information.

### Mice

C57BL/6J wild-type male mice were obtained from Charles River Laboratories (Currie, UK), and male C57BL/6 *mito-*QC mice (constitutive knock-in of mCherry-GFP-mtFIS1^101–152^) were from MRC Harwell Institute (Harwell Campus, Oxfordshire, UK). Under steady-state conditions, the mitochondrial network exhibits both mCherry and GFP fluorescence, whereas during mitophagy, GFP fluorescence is quenched by the acidic microenvironment of the lysosome, leaving punctate mCherry-only foci representing mitolysosomes as an index of mitophagy.^16^ CKD-MBD was induced by feeding a casein-based diet containing 0.6% calcium, 0.9% phosphate and 0.2% adenine (Envigo, Teklad Co. Ltd, Madison, USA). Control (CTL) mice received the same casein-based diet without adenine.^17^ For the RNA-seq study, forty 8-week-old C57BL/6J wild-type mice were randomly allocated in groups of 4 to the control group (n = 12) and sacrificed after 0 (D0), 20 (D20), and 35 (D35) days or the CKD-MBD group (n = 28) and sacrificed every 5 days from day 5 (D5) - day 35 (D35) (Figure 1a). This experimental design assumed little variation in gene expression by bones of young adult CTL mice over 5 weeks, which in line with the 3Rs allowed us to minimise the number of mice studied. In a second study, CKD-MBD was generated in 8-week-old male *mito*-QC mice maintained on the adenine-supplemented (n = 4) or control diet (n = 4) for 5 weeks. At 13 weeks of age, all mice were sacrificed, and blood was obtained by cardiac puncture under terminal anaesthesia. Femurs, tibias and kidneys were harvested, processed, and stored accordingly.

**Figure 1.**
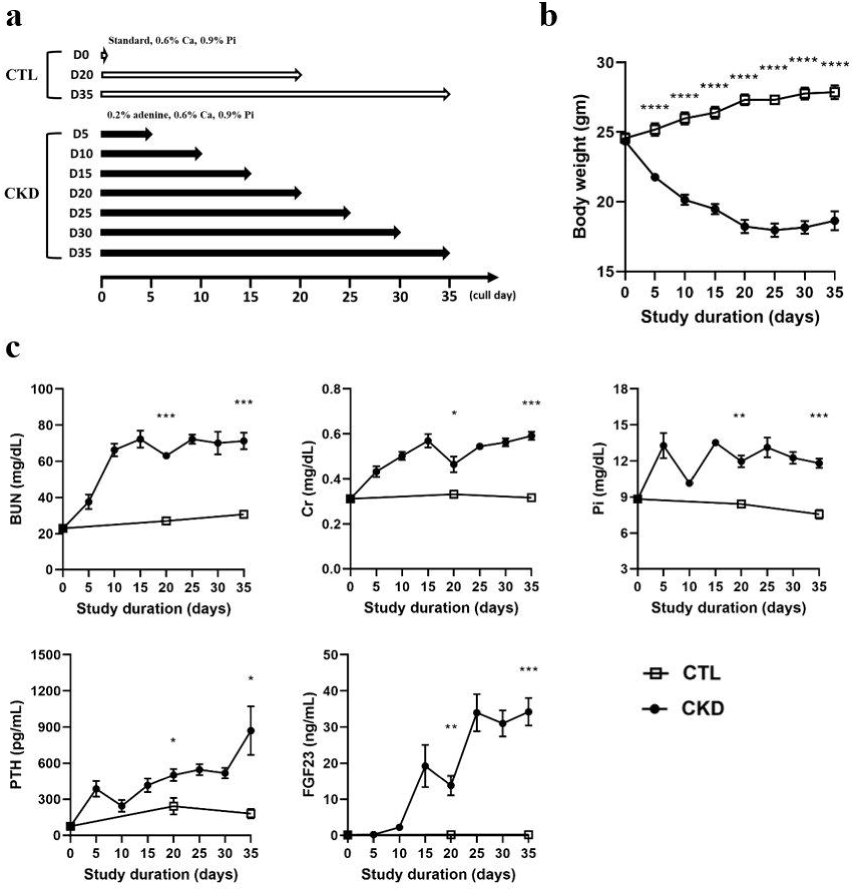
Experimental design for RNA-seq study and time-dependent changes in body weight and serum biochemistries. (a) 40 x 8-week-old male C57BL/6J mice were randomly allocated into 10 groups of 4 mice to the CTL group (n = 12) and sacrificed after D0, D20, and D35 or the CKD-MBD group (n = 28) and sacrificed every five days from D5 to D35. (b) Bodyweight progressively decreased in the CKD-MBD mice, which began losing weight by D5, plateauing at ∼D25. (c) Serum levels of BUN, Cr, Pi, PTH, and FGF23 were measured in mice sacrificed at each sampling point. The data are represented as mean ± SEM (n = 4). NS, not significant, * *p* < 0.05, ** *p* < 0.01, *** *p* < 0.001, **** *p* < 0.0001 vs. CTL mice of the same age.

### Human femur samples

Femoral heads were collected from CKD (n = 3) and non-CKD (n =3) patients going through hip fracture surgery and cortical bone protein samples were prepared for immunoblotting.

### RNA-seq analysis

Barcoded RNA-seq libraries were prepared by Edinburgh Genomics using the Illumina TruSeq stranded mRNA-seq kit (Illumina, Cambridge, UK). RNA (> 1200 ng) from tibial cortical bone was chemically fragmented to yield segments, reverse transcribed using random hexamers, and ligated to barcoded adapters. Sequencing was performed to a depth of 20.6 - 35.5 million mapped paired reads per sample. See Supplementary data for details of mRNA library preparation and downstream network co-expression and functional enrichment analysis.

### Tissue analysis

#### Blood

Serum was analysed for markers of CKD-MBD and bone turnover markers *Micro-computed tomography (*μ*CT):* The cortical geometry and trabecular architecture of the tibia were determined by μCT (Skyscan 1172 X-ray microtomography, Bruker, Kontich, Belgium).

#### Visualisation of actin filaments and mitophagy in mito-QC mice

Cryosections were stained with Alexa Fluor 647 phalloidin and imaged by confocal microscopy. Paraffin sections were imaged directly for the detection of mitophagy.

#### Immunoblotting

Protein from femoral diaphyseal bone (mouse) and femoral head (human) and primary osteoblast cultures was extracted and immunoblotted following standard procedures.

### Osteoblast cell culture

#### Osteoblast isolation

Calvarial osteoblasts were isolated from 3 to 5-day-old C57BL/6J wild-type pups and expanded in flasks for experimentation with indoxyl sulfate (IS) and rapamycin.

#### Mitochondrial staining and determination of membrane potential (ΔΨ) and reactive oxygen species (ROS)

Mitotracker Red stained cells were imaged by confocal microscopy and quantified by spectrophotometry. Cell and mitochondria ROS production was quantified by fluorescence spectrophotometry.

#### Measurement of respiration and glycolysis

This was done using Seahorse XF technology (Agilent Technologies LDA UK Ltd)

#### Immunofluorescence of mitophagy and autophagy associated proteins in primary osteoblasts

Indirect immunofluorescence was used to detect PARKIN, TOM20, LC3 and p62/SQSTM1 using AlexaFluor-conjugated secondary antibodies.

### Statistics

Data are expressed as mean ± standard error of the mean (S.E.M) of at least three biological replicates per experiment. The precise number (n) is indicated in the relevant table and figure legends. Statistical analysis was performed using a two-tailed Student’s t-test or one-way analysis of variance (ANOVA) followed by Tukey’s range test, as appropriate. Statistical analysis was implemented by GraphPad Prism software (GraphPad Software, Inc., USA). *p* values of < 0.05 (*), < 0.01 (**), < 0.001 (***), and < 0.0001 (****) were considered significant.

## RESULTS

### Dietary adenine-supplemented mice develop a renal osteodystrophy phenotype

We first confirmed that mice fed an adenine-supplemented diet presented with a serum profile and skeletal phenotype typical of CKD-MBD and ROD, respectively. CKD-MBD mice began losing weight by D5 and developed a ∼30% reduction of body weight by D35 (Figure 1b, all *p* < 0.0001). Notably, CKD-MBD mice presented with hyperphosphatemia, hyperparathyroidism, and increased serum BUN, Cr, and FGF23 levels (Figure 1c, all *p* < 0.05) relative to CTL mice. This altered serum profile of mice offered an adenine-supplemented diet corresponds in severity to an advanced stage of human CKD.^18^ The trabecular bone microarchitecture, cortical bone geometry and biomechanical properties of the tibia from CKD-MBD mice were also compromised and presented with a typical ROD phenotype (Figure S1a-d). Having confirmed this, we next completed an RNA-seq analysis to identify changes to the bone transcriptome during the development of ROD.

### Experimental ROD is associated with clusters of differentially expressed genes

A Pearson correlation (r ≥ 0.98) matrix was constructed to define relationships between samples using Graphia,^19^ the network analysis program. The 3D visualisation (Figure 2a) of the correlation between the individual sample groups identified CTL samples (n = 12, green dots) in the periphery and CKD-MBD samples (n = 28, blue dots) in a more central position within the major cluster. Unexpectedly, CKD-MBD samples, which included samples collected between D5 and D35, were less heterogeneous than CTL samples (D0, D20, and D35), suggesting reduced bone cell activity. Principal component analyses (PCA) disclosed that the first two principal components (PC1 + PC2) explained 56.1% of the data variance (Figure 2b).

**Figure 2.**
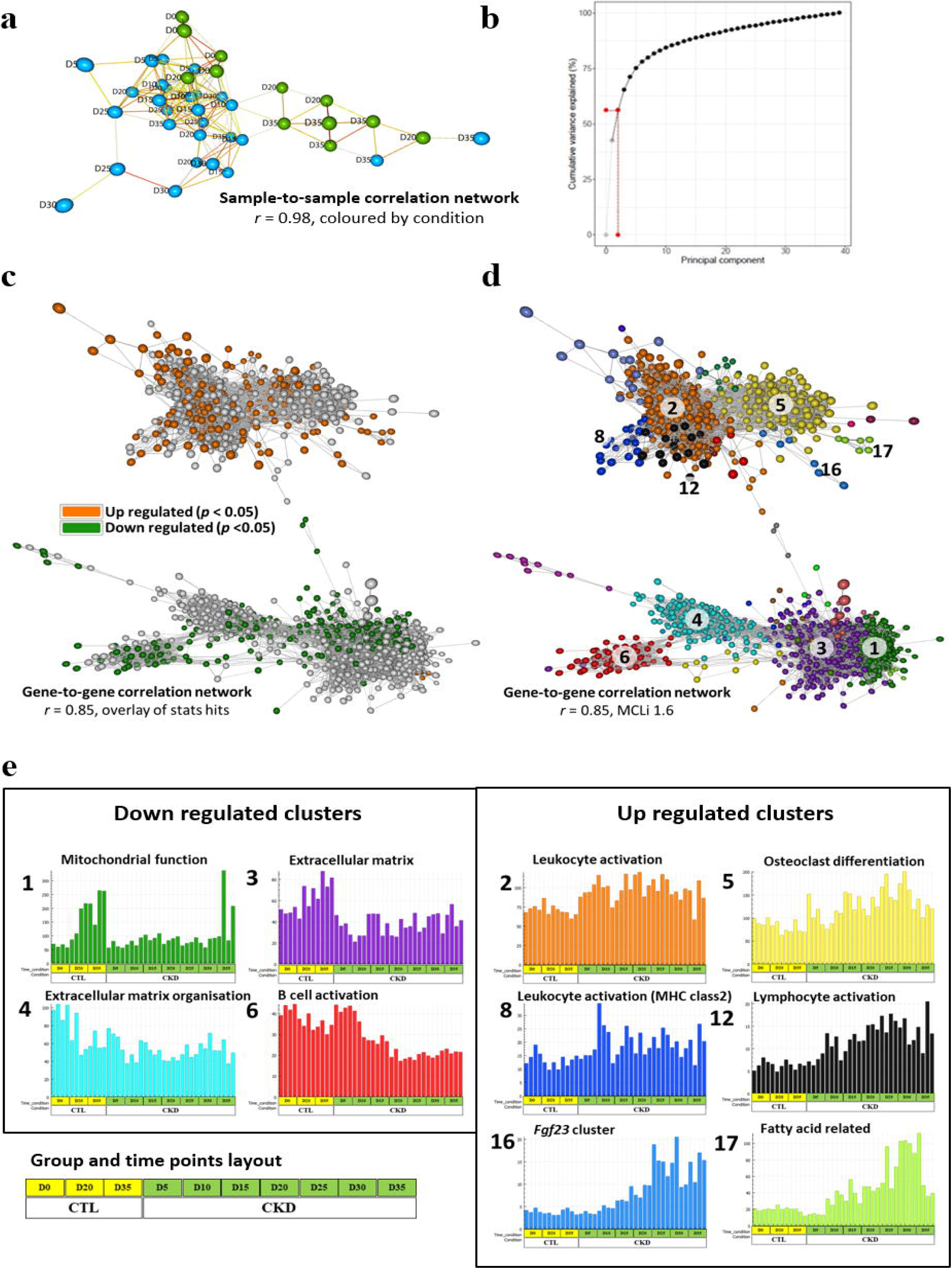
Network visualisation and analysis reveal CKD-MBD correlated gene clusters with bespoke biological functions. (a) Three-dimensional visualisation of a sample-to-sample Pearson correlation network of control samples (n = 12, green nodes) and CKD samples (n = 28, blue nodes) showed late time points to be generally more distinct from early time points, and for the most part control samples to different from CKD samples. The strength of edges (line) was represented on a scale from thin and blue (weak) to thick and red (strong). It corresponded to correlations between individual measurements above the defined threshold (r ≥ 0.98). (b) The red line indicated that the first two principal components (PC1 + PC2) explain 56.1% of the variance of the data. (c) A co-expression graph of the RNA-seq data was constructed comprised of 3,455 nodes (genes) and 730,900 edges (correlations r ≥ 0.85). The graph was made up of two large and unconnected components of genes representing those whose expression was higher or lower in the CKD samples relative to controls as seen when differentially expressed genes with ≥ 1.5-fold change from D20 and D35 samples were overlaid on the network. The group highlighted in orange contains 411 transcripts whose expression was significantly up-regulated following adenine-induced CKD-MBD at D20 and D35. The group highlighted in green includes 754 transcripts consistently down-regulated following adenine-induced CKD-MBD at D20 and D35. (d) The network’s nodes represent CKD-MBD correlated genes, and edges represent correlations above a defined threshold (r ≥ 0.85). Co-expressed genes form highly connected complex clusters within the graph. A gene-to-gene correlation graph was generated from all genes that were correlated (r ≥ 0.85) with differentially expressed genes from D20 and D35 (CKD vs. control). Nodes were coloured by cluster using the MCL algorithm (1.6) according to their specific biological function. The edges corresponded to the correlation between them. (e) Ten histograms showing the mean expression profile of gene clusters. Further functional enrichment analysis identified the biological processes associated with the genes in each of these 10 clusters. Those associated with *Fgf23* function, osteoclast differentiation, fatty acid synthesis, and leukocyte and lymphocyte activation increased with time, whereas genes associated with mitochondria function, B cell activation, and extracellular matrix structure and organisation decreased over time.

A gene-to-gene correlation network was generated with a Pearson correlation threshold (r ≥ 0.85) and a Markov clustering (MCL) inflation value of 1.6 was used to define relationships between genes. The correlation network comprised nodes (genes) correlated with CKD-MBD after enrichment analysis and the formation of clusters of differentially expressed genes from D20 and D35 samples (Figure 2c). The network included 3,455 nodes and 730,900 edges (links). Two distinct groups of nodes sharing the same correlation factor were selected for functional analysis. The top representative group (≥ 1.5-fold up-regulated, highlighted in orange) contained 411 transcripts whose expression was up-regulated in bones from CKD-MBD mice. In contrast, the bottom group (≤ 1.5-fold down-regulated, highlighted in green) had 754 transcripts consistently downregulated in bones from CKD-MBD mice. All other genes (highlighted in grey) identified within both groups were highly correlated and consistently down- or up-regulated across all time points (D5 to D35) of CKD-MBD mice but did not reach the 1.5-fold threshold. The smallest cluster size of the two-node algorithm was used to subdivide the network into discrete gene clusters that contained genes with similar biological functions and expression patterns over time. For example, the expression profiles of the five genes in the *Fgf23* cluster (cluster 16) are shown in Figure 2d and Figure S2.

### The expression of genes associated with mitochondrial function is altered in renal osteodystrophy

The resultant correlation network comprised 41 gene clusters related to the development of ROD; there were seventeen clusters of up-regulated genes and twenty-four clusters of down-regulated genes (Figure 2d). Genes found in each cluster are listed in supplementary data file 1, and clusters 1 to 41 (numbered in order of size; *e.g.,* cluster 1 comprised 1675 genes) were annotated. Of the 41 clusters, ten major clusters possessed biological functions associated with bone metabolism/(re)modelling, and additional functional enrichment analysis using ToppGene identified the biological process associated with the genes in each of these 10 clusters (Figure 2e). Clusters 2, 8, and 12 indicated that activation of leukocytes and lymphocytes accompanied ROD development, whereas cluster 6 revealed the suppression of B cell function. These data confirm previous observations that CKD is a chronic inflammatory state.^20^ Increased osteoclast differentiation (cluster 5) and altered extracellular matrix structure and organisation (clusters 3 and 4) were consistent with increased bone (re)modelling during the onset and development of ROD.^21–23^ Similarly, increased bone marrow fat accumulation and *Fgf23* expression are recognised hallmarks of CKD-MBD.^24, 25^ An unexpected finding was the downregulation of genes associated with mitochondrial function and energy metabolism (cluster 1) during the onset and development of ROD (Figure 2e).

### Identification of mitochondrial pathways downregulated in renal osteodystrophy

To identify mitochondrial and energy-related pathways implicated in the aetiology of ROD, we next performed WebGestalt enrichment analyses on the genes that comprise cluster 1 (Figure 3a). The top 10 gene ontology (GO) terms identified included the tricarboxylic acid (TCA) cycle, oxidative phosphorylation, and neurodegenerative disorder-associated pathways such as Parkinson’s and Alzheimer’s disease. Specifically, heat-maps identified genes in cluster 1 that were down-regulated in pathways associated with glycolysis and mitochondrial function, such as the TCA cycle, mitophagy (the selective degradation of damaged mitochondria by autophagy), and mitochondrial ETC complexes I to IV (Figure 3b). The expression of mitophagy-associated genes in CKD-MBD samples did not, unlike CTL samples, increase over the experimental period.

**Figure 3.**
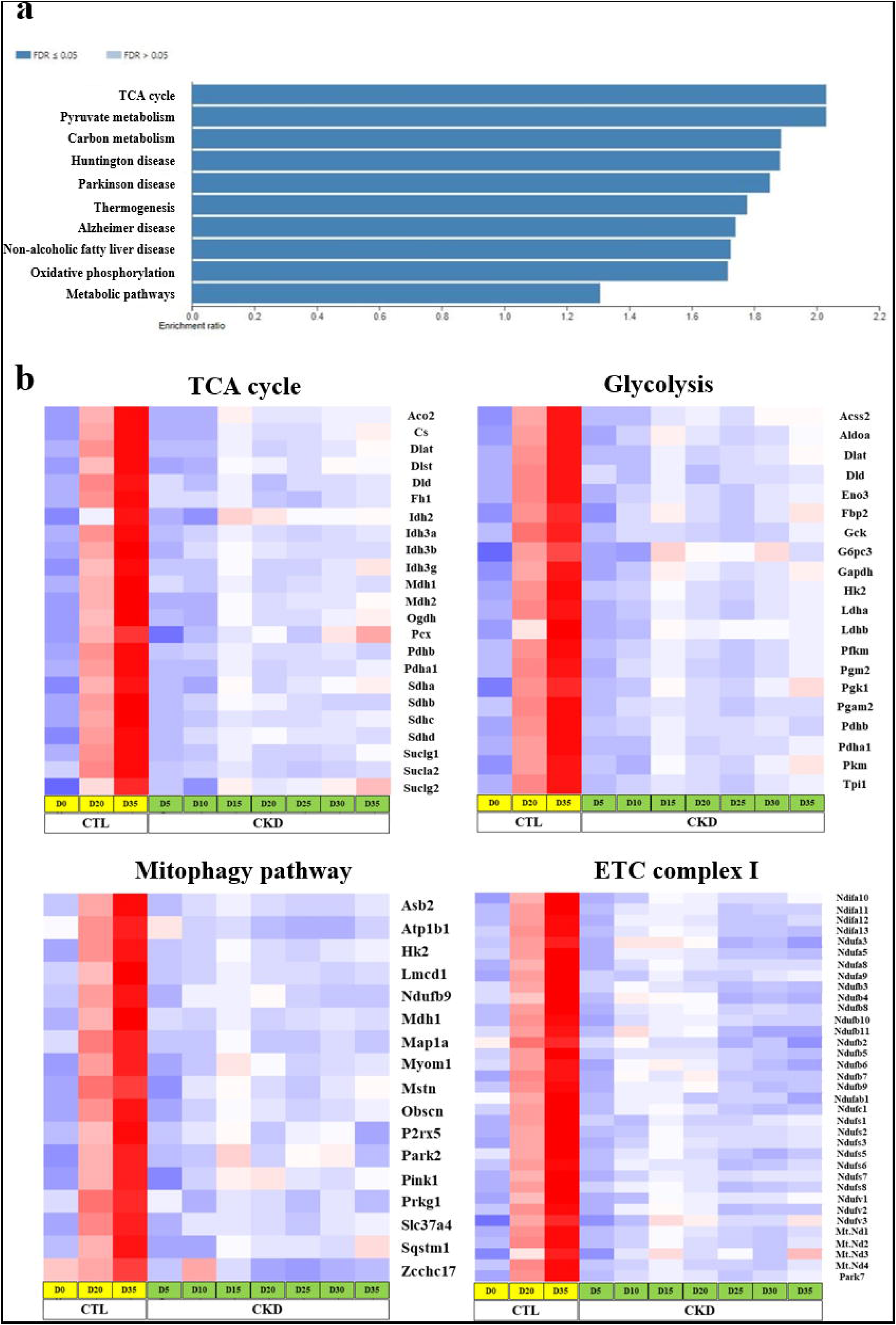
Pathway analysis of differentially expressed genes in Cluster 1 of the RNA-seq data. (a) The top 10 GO terms are listed. (b) Heatmaps were derived from the average value of four biological replicates at each time point and showed that genes associated with the TCA cycle, glycolysis, mitophagy, ETC complex I [and II, III, and IV (not shown)], were down-regulated in CKD-MBD mouse bones.

### Mitolysosomes and p62 puncta are increased in osteocytes from renal osteodystrophy mice

We speculated that a reduced clearance of damaged mitochondria detected in experimental ROD could underpin the down-regulation of energy production pathways in mitochondria.^26^ To explore this further and quantify mitophagy within osteocytes (the most abundant cell type in bone and the likely source of the RNA-seq transcriptome data), we induced CKD-MBD in *mito*-QC mice. There was a 2.3-fold increase in red puncta (mitolysosomes) in osteocytes situated within the cortical bone of CKD-MBD mice (Figure 4a-c). The accumulation of osteocyte mitolysosomes in ROD could indicate either enhanced mitophagy activation or a blockage of downstream steps in the process of mitochondria removal. The accumulation of damaged mitochondria is more likely as this would be more harmful to bone than increased mitophagy. Interestingly, in bone of CKD-MBD mice, the dendritic processes radiating from the osteocyte cell body were reduced in number (Figure S3). These reduced dendritic processes may impede the endoplasmic reticulum mediated removal of defective osteocyte mitochondria in ROD.^27^ Of note, increased mitolysosomes in CKD-MBD were not specific to bone as they were also observed in the renal tubules and glomeruli of CKD-MBD mice (Figure S4).

**Figure 4.**
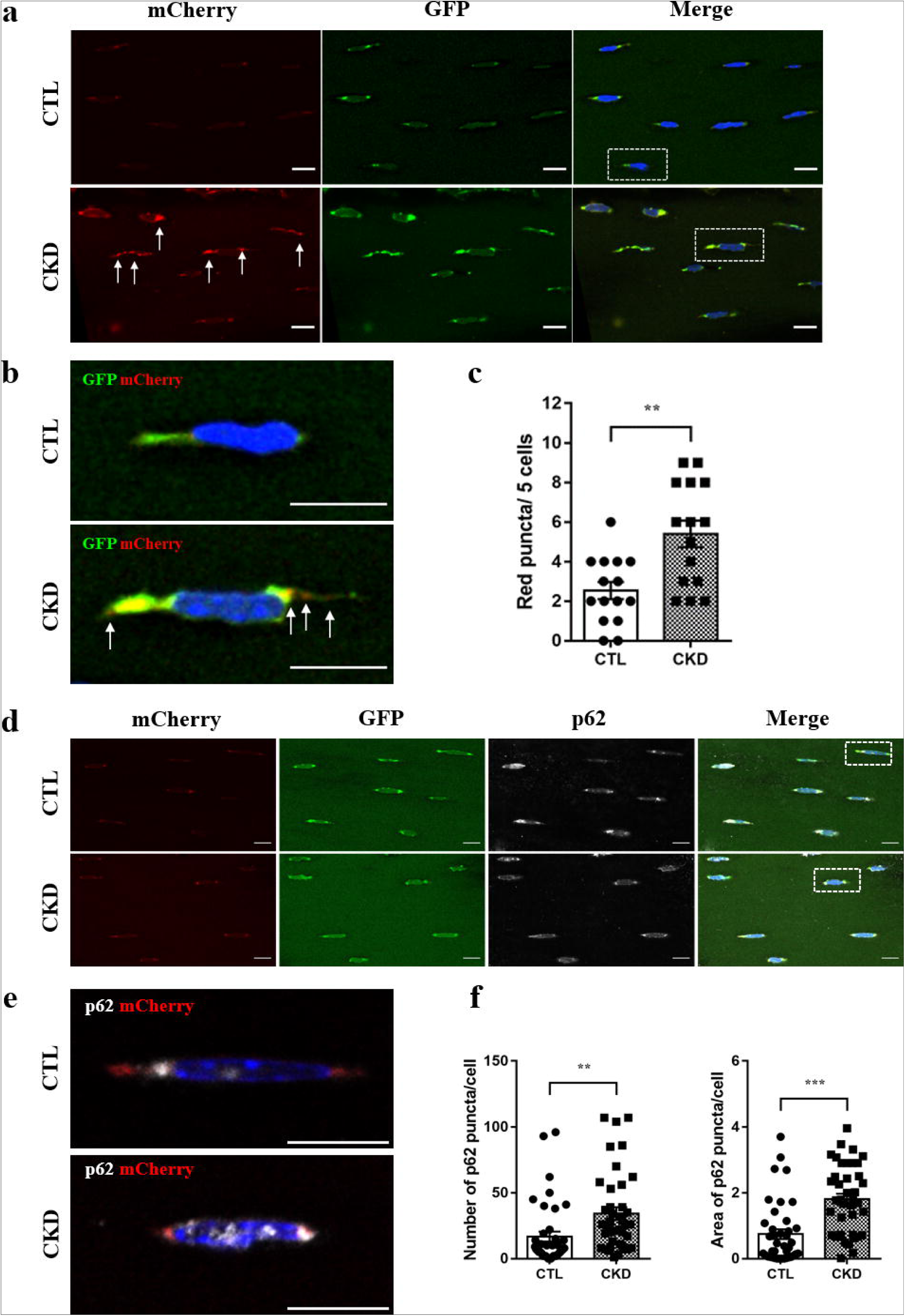
Quantification of mitophagy in osteocytes from CTL and CKD-MBD *mito*-QC mice. (a) Cortical bones from CTL and CKD-MBD *mito*-QC mice were imaged, whereby significant red mitophagic puncta were identified in osteocytes of CKD-MBD bones (white arrowheads). (b) Red mitophagic puncta (white arrowhead) were observed at higher magnification within the osteocyte cytoplasm and emerging dendrites. (c) Quantification of red mitophagic puncta is shown where each dot represents the total number of red puncta/5 osteocytes/section. (d) Assessment of the selective autophagy adaptor p62/SQSTM1 in CTL and CKD *mito*-QC mice osteocytes. Cortical bones from CTL and CKD *mito*-QC mice were imaged, whereby p62/SQSTM1 were identified in osteocytes of CKD bones. (e) Representative images of white p62 co-localised with red mitolysosomes showing abundant p62 accumulation in CKD osteocytes. (f) The average number and area (µm^2^) of p62 co-localised puncta per osteocyte were quantified in CTL and CKD bones. Each dot represents the average number and surface area of the co-localised puncta per cell. Scale bar 10 μm. Data are presented as mean ± S.E.M (5 sections/mouse, n = 3). Analysis was tested by a Student’s t-test. ** *p* < 0.01, *** *p* < 0.001 vs. the CTL group.

### The expression of mitophagy regulatory proteins in osteocytes from CKD-MBD bone suggests a block in the clearance of damaged mitochondria

To identify the mechanisms leading to mitolysosome accumulation in ROD, the expression of established mitophagy regulators was analysed. The expression of PARKIN, an E3 ubiquitin ligase that promotes mitophagy by ubiquitinylation of substrates at the outer mitochondrial membrane was increased in cortical bone from CKD-MBD mice (Figure 5a, b). The autophagic adaptor protein, p62 (sequestosome 1/SQSTM1) is a major LC3-II interacting protein and as a selective substrate it is normally degraded during autophagy.^28^ However, no decrease in p62/SQSTM1 expression by western blotting was noted in bone from CKD-MBD mice, whereas by immunostaining the number of p62/SQSTM1 puncta/ osteocyte was increased in osteocytes of *mito*-QC CKD-MBD mice (Figures 4d-f). Also, the autophagy-related 7 (ATG7) protein, which increases during autophagy and drives the pivotal stages of classical degradative autophagy through the lipidation of LC3-I to LC3-II, was similarly expressed in bone from CTL and CKD-MBD mice (Figure 5a, b). These latter observations are incongruous with increased mitophagy and suggest a block in the removal of damaged mitochondria via the autophagic machinery. An increase in osteocyte mitochondria number was confirmed using the mitochondrial marker TOM20, a subunit of the mitochondrial translocase of the outer membrane complex, supporting the concept that mitochondria are not being cleared in ROD (Figure 5a, b). This was replicated in the cortical bone of CKD-MBD patients, whereas a decrease in ATG7 protein levels was observed (Figure 5c, d).

**Figure 5.**
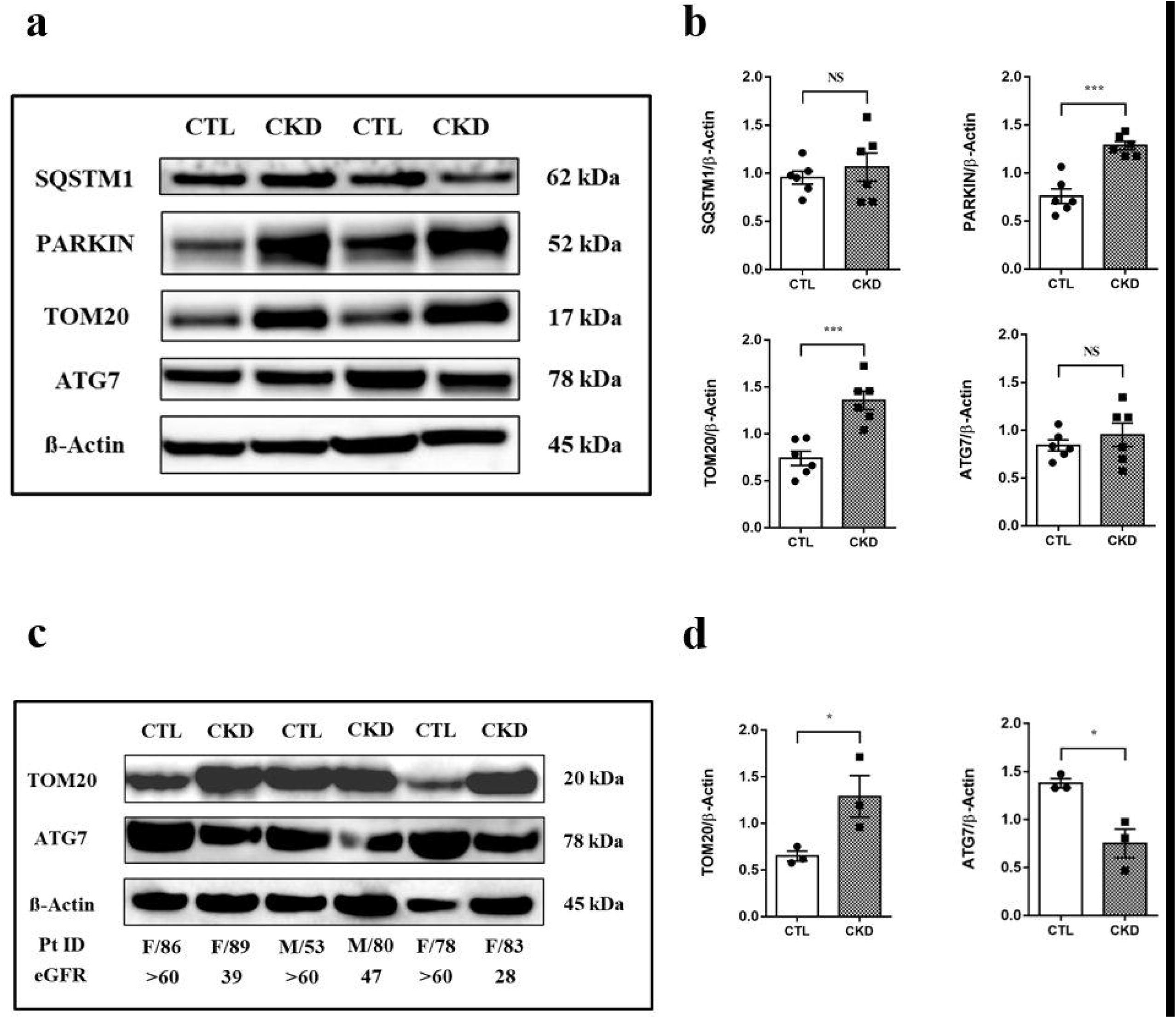
Mitophagy regulators are altered in the femurs of CKD-MBD mice and patients. (a) Representative western blots of two CTL and two CKD.MBD samples showing protein expression levels of mitophagy-associated proteins in cortical bone of mouse femurs. (b) Quantification of SQSTM1, PARKIN, TOM20, and ATG7 from CTL and CKD-MBD mouse femurs in cortical bone. (c & d) Representative western blot analyses and quantification of TOM20 and ATG7 in cortical bone of human femurs. Data are presented as means ± SEM, n = 6 (mouse femurs) and n = 3 (human femurs). Analysis was tested by Student’s t-test. NS, not significant, * *p* < 0.05 and *** *p* < 0.001 vs. the CTL group.

### Indoxyl sulfate (IS) impairs mitochondria function and morphology, and mitophagy in primary osteoblasts

Uremic toxins accumulate in body fluids during progressive CKD and previous studies have reported that they may induce mitochondrial damage and also impair autophagy in various cell types.^29, 30^ Therefore, we hypothesised that the failure to remove damaged mitochondria via impaired mitophagy (Figure 5a-d) may result from increased uremic toxin levels in CKD-MBD mice. To examine this further, we treated murine primary osteoblasts with the uremic toxin, IS (0-2 mM) for 7 days, a treatment regimen that was not toxic to cells (Figure S5). Osteoblast mitochondrial ΔΨ was decreased by IS as evidenced by a reduction in mitotracker red uptake, which is dependent on ΔΨ, whereas IS treatment increased mitochondrial and cellular ROS production in a concentration-dependent manner (Figure 6a-c). The morphology and distribution of osteoblast mitochondria were markedly affected by IS treatment. In CTL cells, the mitochondrial network appeared as long thread-like tubular structures distributed throughout the cytoplasm (Figure 6d, left panel, arrows), which contrasted with the IS-treated cells where the mitochondrial network was disintegrated and characterised by fragmented mitochondria with a swollen rounded morphology, collapsed around the nuclei (Figure 6d, right panel, arrowheads). Determination of mitochondrial oxygen consumption rate (OCR) and extracellular acidification rate (ECAR) in control and IS-treated primary osteoblasts revealed that basal and maximum respiration and glycolysis capacity were decreased by IS, as was ATP production, max H^+^ leak and non-respiratory OCR (Figure 7a-c). These observations are in agreement with the *in vivo* RNA-seq pathway analysis shown in Figure 3b. Finally, the expression of p62**/**SQSTM1 was up-regulated by IS treatment in a concentration-dependent manner, whereas the expression of PARKIN and autophagic marker LC3-II (generated in an ATG7-dependent manner) were downregulated by IS treatment (Figure 8a, b, and Figure S6 & S7).

**Figure 6.**
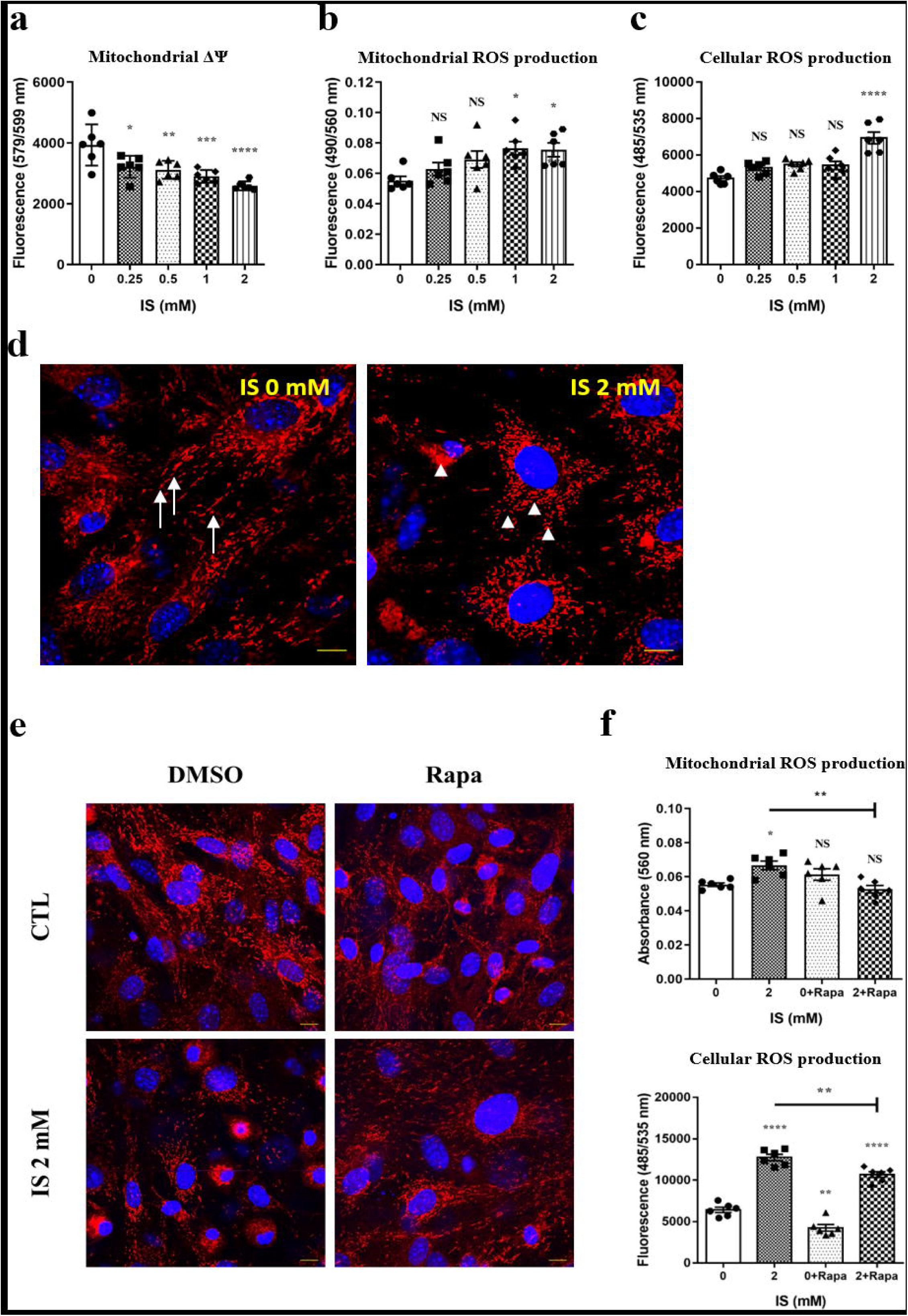
Rapamycin reduces ROS production and rescues IS-induced mitochondrial structural abnormality in IS-treated primary osteoblasts. (a) Quantification of mitochondria ΔΨ in indoxyl sulfate (IS) treated osteoblasts (b) Quantification of mitochondrial and (c) cellular ROS production by osteoblasts treated with varying concentrations of IS. (d) Representative confocal images of primary osteoblasts stained with MitoTracker Red CMXRos. In control cultures, the mitochondrial network appeared as long thread-like tubular structures (left panel, arrows), whereas those treated with IS (2mM) had a swollen, rounded morphology (right panel, arrowheads). (e) Abnormally shaped mitochondria in IS-treated cells (2 mM) were restored by adding rapamycin (75 nM). (f) Quantification of mitochondrial and cellular ROS production by osteoblasts treated with 0 or 2mM indoxyl IS ± rapamycin. Scale bar, 10 μm. The data are represented as the mean ± SEM (n = 6), NS, not significant, * *p* < 0.05; ** *p* < 0.01; *** *p* < 0.001; **** *p* < 0.0001 vs. non-treated CTL group (0 mM).

**Figure 7.**
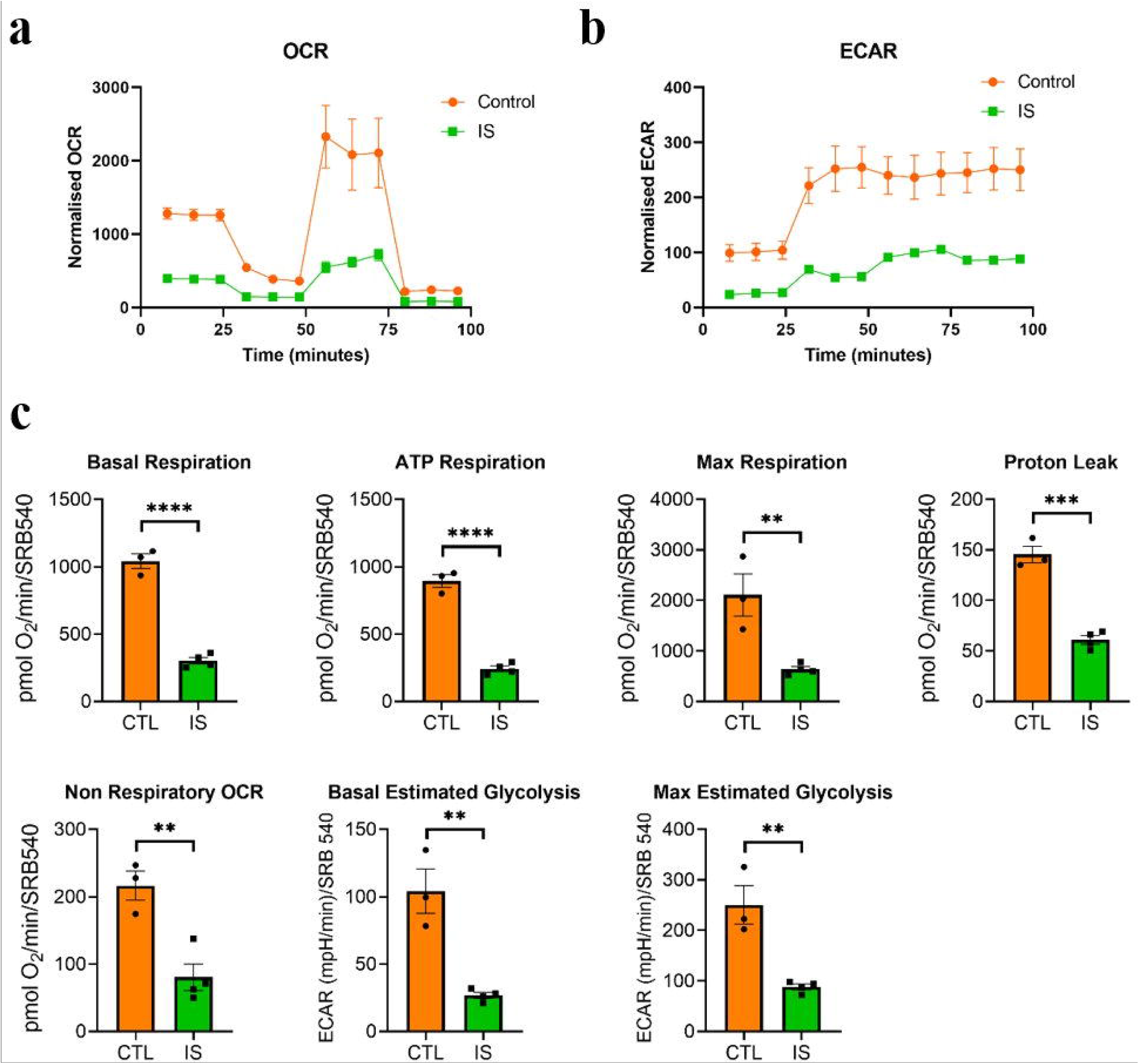
The effects of IS on oxidative phosphorylation and glycolysis in primary osteoblasts. Cells were treated in the presence/absence of indoxyl sulfate (IS; 0 vs. 2 mM) for 7 days. (a & b) Mitochondrial oxygen consumption rate (OCR) and extracellular acidification rate (ECAR) in primary osteoblasts following the addition of oligomycin (oligo), carbonyl cyanide 4-(trifluoromethoxy) phenylhydrazone (FCCP) and rotenone/antimycin A (AR1) to target the electron transport chain. 1.5 μM FCCP for the controls was used as optimal, and 3 μM FCCP for IS as optimal. (c) From the OCR and ECAR data, basal respiration, ATP-linked respiration, maximal respiration, proton leak, non-respiratory OCR, basal estimated glycolysis, and maximal estimated glycolysis were calculated. The data are represented as the mean ± SEM (n = 3), ** *p* < 0.01; *** *p* < 0.001; **** *p* < 0.0001 vs. non-treated CTL group (0 mM).

**Figure 8.**
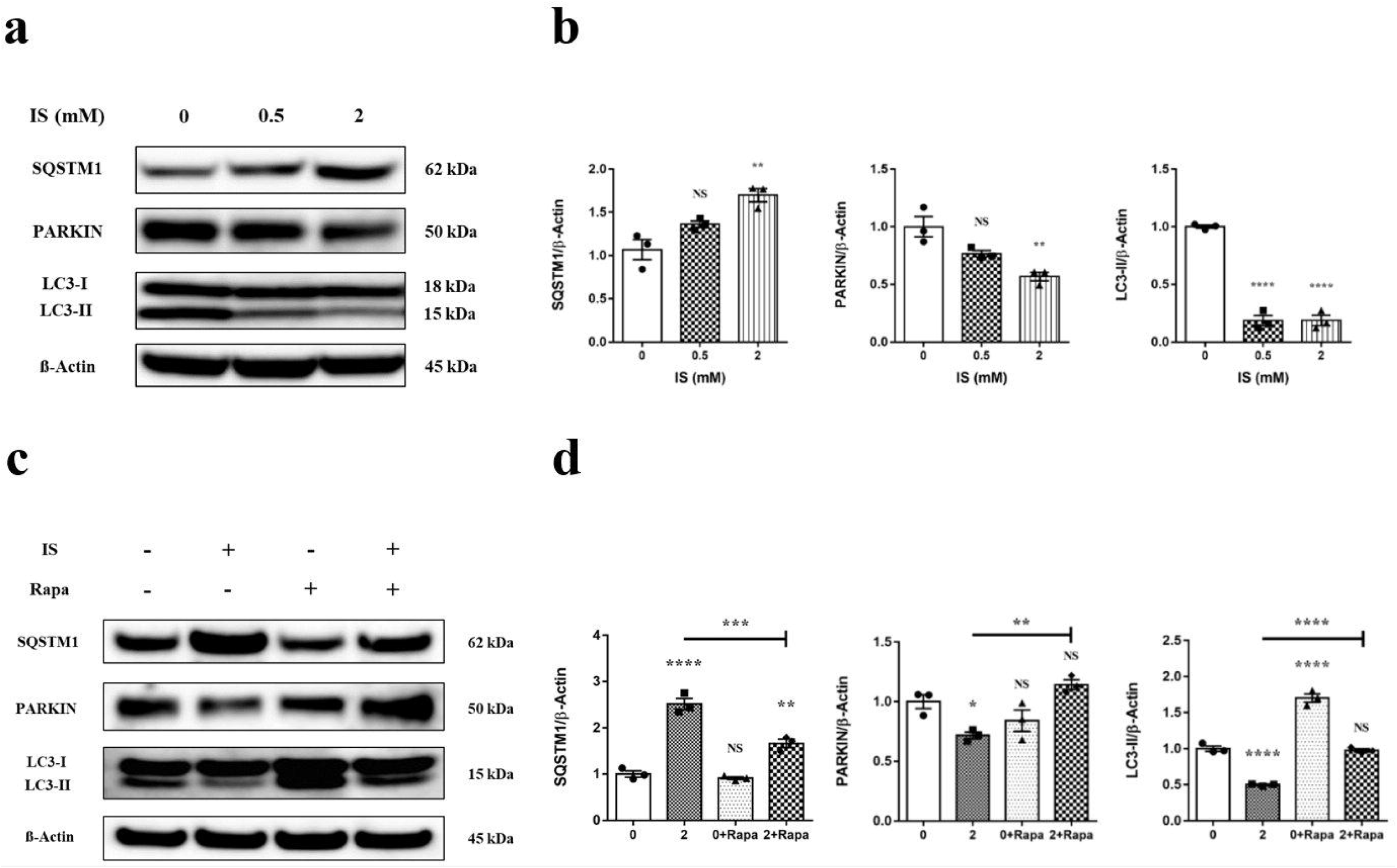
Rapamycin reverses the adverse effects of IS on primary osteoblast mitophagy. The expression and quantification of mitophagy regulators by primary osteoblasts treated with increasing concentrations of indoxyl sulfate (IS) (0, 0.5, and 2 mM) for 7 days. (a & b) The effect of IS (0 - 2 mM) on the expression of PARKIN, SQSTM1, and LC3-II. (c & d) The ability of rapamycin to reverse the effects of IS on the expression of SQSTM1, PARKIN, and LC3-II. The signal intensity was normalised to β-Actin and set at 1 for the control cultures (no IS) (n = 3). Significance was calculated by one-way ANOVA with a post hoc Tukey’s multiple comparisons tests. NS, not significant, * *p* < 0.05; ** *p* < 0.01; **** *p* < 0.0001 vs. untreated CTL group (0 mM).

### Rapamycin rescues primary osteoblasts from the adverse effects of IS on mitochondrial morphology and clearance, ROS production and mitophagy

Rapamycin, a specific mammalian target of rapamycin (mTOR) inhibitor, is a commonly used autophagy inducer that increases oxidative phosphorylation, decreases ROS production, and, by removing damaged/dysfunctional mitochondria it improves overall mitochondrial quality.^31, 32^ Cells (± IS; 2 mM) were treated with vehicle (control) or rapamycin (± Rapa; 75 nM) for 7 days, and the expression of mitophagy-associated proteins (PARKIN, LC3-II, and p62/SQSTM1) was determined. The reduction of PARKIN expression by IS was reversed by rapamycin, resulting in an increase in PARKIN expression in the IS + rapamycin-treated cells (Figure 8c, d, and Figure S6). Also, rapamycin (with or without IS) increased LC3-II expression and reversed the reduced conversion of LC3-I to LC3-II in IS-only treated cells (Figure 8c, d). Finally, rapamycin also partly reversed the IS-increased p62/SQSTM1 expression (Figure 8c, d and Figure S7). We next determined whether the promotion of mitophagy by rapamycin could prevent or mitigate IS-induced mitochondrial structural damage and dysfunction. The oxidative stress induced by IS was lowered by co-incubation with rapamycin. Furthermore, the ability of rapamycin to reactivate IS-suppressed mitophagy and efficiently remove damaged mitochondria resulted in the correction of the abnormal morphology and distribution of mitochondria (Figure 6e-f). These results imply that rapamycin was able to rescue cultured osteoblasts from the adverse effects of IS on mitochondria morphology and mitophagy removal.

To further investigate the role of LC3 in rapamycin’s ability to promote mitophagy and rescue the cells from IS challenge, we treated osteoblasts with trifluoromethoxy carbonylcyanide phenylhydrazone (FCCP), a potent mitochondrial uncoupling agent that dissipates mitochondrial membrane potential and triggers mitophagy by recruiting PARKIN to damaged mitochondria.^33, 34^ Following IS challenge, there was increased mitochondrial damage but fewer LC3 puncta cell. Rapamycin exposure enhanced LC3 recruitment to mitochondria (Figure 9a, b, yellow puncta) in both control and IS-treated cells in the presence of FCCP. We hypothesise that FCCP may enhance the clearance of damaged mitochondria by triggering the expression of mitophagy signals (*e.g.,* PARKIN) thus, recruiting rapamycin-induced LC3-II to them (Figure 9b).

**Figure 9.**
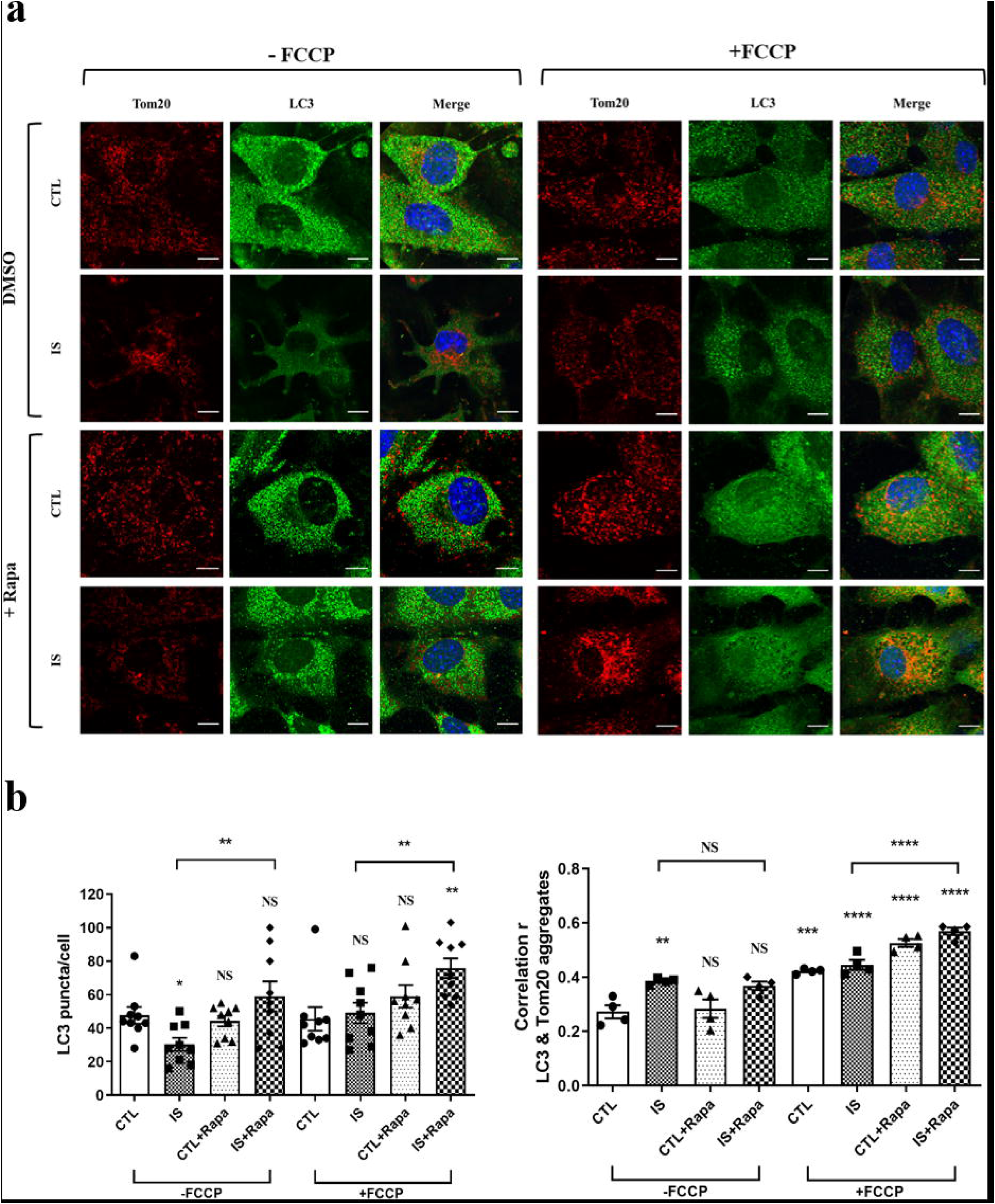
Rapamycin stimulates mitophagy by increasing LC3 puncta that co-localises with TOM20 in IS-treated primary osteoblasts. (a & b) Quantification of LC3 puncta and its co-localisation with the outer mitochondrial marker TOM20 in the presence or absence of indoxyl sulfate (IS), rapamycin, and carbonyl cyanide 4-(trifluoromethoxy) phenylhydrazone (FCCP). Autophagic flux (puncta) is accurately measured via the inhibition of lysosomal degradation by bafilomycin treatment for 6 hrs, followed by cell fixation. The number of LC3 puncta was counted in 9 osteoblasts from at least three images per treatment group. Scale bar = 10 μm. The data are represented as the mean ± SEM, NS, not significant, * *p* < 0.05; **** *p* < 0.0001 vs. vehicle-DMSO CTL group in the absence of FCCP.

## DISCUSSION

It is well-established that CKD-MBD is associated with poor bone health, but many aspects of the pathophysiology remain unclear. Therefore, to obtain an improved understanding of the biological processes disrupted during the initiation and development of ROD, we completed an initial RNA-seq study on cortical bone from a rodent model of CKD-MBD. This approach identified a number of pathways and functions associated with bone metabolism/remodelling that were altered in the CKD-MBD mice. These included disordered osteoclast resorption and extracellular matrix structure and organisation, which are recognised to be involved in the aetiology of ROD.^17, 21, 22^ However, altered osteoblast/osteocyte mitochondrial function has not previously been reported and warranted further study.

This is a novel observation in bone and is in keeping with previous reports in which mitochondria dysfunction has been associated with pre-CKD kidney diseases.^35^ Oxidative stress may be involved, but due to their many integrated functions in addition to their indispensability for cellular respiration and oxidative phosphorylation, the precise nature of the mitochondrial insult remains obscure.^36^ Whilst the kidney is particularly susceptible to oxidative stress due to its high energy demands, mitochondria are also essential for maintaining bone homeostasis by controlling the balance between osteogenesis and bone resorption.^37^ Altered mitochondrial function, *e.g.,* mitophagy, fusion, fission, and biogenesis, is associated with decreased bone mass, and mitophagy-focused studies have reported that irregular mitophagy via decreased PINK or PARKIN leads to impaired osteogenic differentiation.^38–40^ Impaired autophagic flux noted in osteoblasts and osteocytes of ATG7 null mice may explain the observed lower bone mass, reduced remodelling, and fewer osteocyte projections.^41, 42^ Furthermore, accumulating mitochondrial DNA (mtDNA) mutations in mice results in the premature onset of ageing-related phenotypes, including osteoporosis and vertebrae compression fractures.^43–45^ Corroborating clinical data is limited, but mitochondria point mutations, including the common m3234A>G, are associated with decreased bone mass and strength, and severe osteoporosis.^46, 47^ The causative insult(s) are likely to be complex, but oxidative stress stimulates osteoclast formation and inhibits osteoblast differentiation via increased sclerostin and the downregulation of Wnt/β-catenin signalling.^45, 48, 49^

From the RNA-seq data of this study, we speculated that impaired mitophagy in CKD-MBD bones would lead to the retention of damaged mitochondria and could explain the decreased oxidative phosphorylation and contribute to the aetiology of ROD. However, the reduced mitophagy as implied by the RNA-seq data, is at odds with the increased PARKIN protein and mitolysosomes noted in the CKD-MBD *mito*-QC mice. An explanation for this apparent paradox is undoubtedly complex but may be based on the pretext that changes in gene function (protein levels) can be extrapolated from changes in transcript levels.^50^ Indeed, in the ageing murine kidney, mRNA transcript levels have been reported to be a poor proxy for protein abundance which may be a consequence of changes in translational efficiency and/or protein turnover.^51^ This phenomenon is not uncommon in CKD and ageing animal studies focused on renal physiology.^52, 53^ This disconnect between transcript and protein levels was further emphasised when the expression of established mitophagy regulators in CKD-MBD bone was quantified by immunoblotting. In CKD-MBD mouse bone, increased TOM20 and the preservation of ATG7 and p62/SQSTM1 levels are inconsistent with their recognised role in the mitophagy process and are indicative of dysfunctional or blocked mitophagy.^54–56^ While the presence of increased red mitolysosomes in the osteocytes of CKD-MBD bone implies increased mitophagy and high baseline turnover, this observation may be misleading due to slower degradation of mitolysosomes at the autophagosome stage. A compromised ability to efficiently and quickly remove damaged mitochondria as a consequence of dysfunctional mitophagy has been reported previously in a number of non-skeletal organs.^57–59^

In an attempt to determine the driver for the altered mitophagy noted in our experimental ROD model, we focused on uremic toxins because of their known ability to damage mitochondria and impair autophagic flux.^29, 30^ The effects of IS on isolated osteoblasts were consistent with aberrant mitophagy (increased p62/SQSTM1 and decreased PARKIN and LC3-II), but the decreased PARKIN protein is at odds with what is noted in CKD-MBD mice (Fig 5). This suggests that the effects of IS alone do not fully mimic the *in vivo* situation where other serum factors altered in CKD-MBD may mitigate the actions of uremic toxins on the mitophagy process. Moreover, although mitochondrial damage was observed in IS-treated cells, there was less LC3 punctate/LC3-II within damaged mitochondria, implying reduced PARKIN to signal the autophagosomes (Figures 9a-b and 8a-b). However, as rapamycin reversed the effects of IS on PARKIN, LC3-II, p62/SQSTM1 and punctate LC3 (Figures 8c-d, and 9a-b, left panel), it raises rapamycin and other mitochondria-targeted approaches as a potential pharmacological intervention to manipulate mitochondrial function within cells and reduce CKD-related bone disorders.^57, 60, 61^ Rapamycin, an inhibitor of mTOR, is a drug clinically used for antifungal, immunosuppressive, and anti-tumour purposes; however, its application to augment mitophogy in the complex pathophysiology of ROD may be limited due to its harmful impact on bone formation.^62^

We have speculated that the increased ROS production by mitochondria in bone cells in response to uremic toxins may underpin the altered mitophagy noted in this study. Uremic toxins accumulate in the tissues and serum of CKD patients and have been implicated in the earlier adynamic bone disease phase of ROD, which is characterised by skeletal PTH resistance.^63, 64^ Uremic toxins inhibit osteoblast differentiation and matrix synthesis and alter bone’s chemical composition, resulting in impaired biomechanical properties and increased fragility.^63, 65, 66^ IS may promote these bone pathologies by deregulating mitochondrial metabolism, mass, and dynamics via the induction of oxidative stress by activating NADPH oxidase.^67, 68^ Disrupted oxidative phosphorylation in IS-treated osteoblasts was noted in this study, as was an accumulation of p62/SQSTM1 and reduced LC3-II protein, which together exemplify impaired mitophagy. Furthermore, AST-120, an oral adsorbent of uremic toxins, has been shown to slow CKD progression, possibly by reducing oxidative stress and suppressing mitochondrial dysfunction.^64, 69^ Clinical studies examining the ability of AST-120 to minimise/delay/prevent ROD have not been reported, but rodent studies have reported that AST-120 can suppress the progression of adynamic bone disease in uremic rats.^63, 70^ These data support the tenet that uremic toxins damage mitochondria through oxidative stress, impact mitophagic function in bone cells and contribute to the ROD phenotype.

This body of work provides persuasive evidence that altered osteoblast/osteocyte mitochondrial function has a major impact on the development of ROD and is consistent with the emerging role of the osteocyte in CKD.^71^ While not the focus of this study, reports indicate that altered mitochondrial function may be central to some of the co-morbidities associated with CKD. Muscle wasting is a severe complication of CKD and probably contributes to the increased risk of falls. Experimental evidence suggests impaired mitochondrial metabolism is a crucial mechanism linking kidney dysfunction with sarcopenia.^72, 73^ Moreover, the protein levels of various mitophagy mediators such as BNIP3, p62, LC3-II, PARKIN and PINK1 are altered in mitochondria isolated from muscle tissues from CKD patients, which may account for the poor muscle mitochondrial oxidative capacity in CKD patients.^57, 74, 75^ Interestingly, L-carnitine and teneligliptin, a dipeptidyl peptidase-4 (DPP-4) inhibitor, normalise muscle mitophagy markers and reduce CKD-induced muscle atrophy in mice.^61^

In summary, we have identified the cellular mechanisms implicated in the aetiology of ROD, using an RNA-seq study on bones from a CKD-MBD mouse model. This and follow-up studies revealed evidence of mitochondrial dysfunction and repressed mitophagy within osteoblasts/osteocytes (Table 1). As mitochondria are critical for osteoblast/osteocyte function, and mitochondrial respiratory chain dysfunction in osteoblasts results in accelerated bone loss, we propose that the altered uremic toxin levels characteristic of CKD-MBD impair mitochondrial function and contribute to the development of ROD.

**Table 1.** Expression of mitophagy regulators in both *in vivo* and *in vitro* models of CKD.

|  | **CKD gene**<br>**(Fig 3)** | **CKD protein**<br>**(Fig 5)** | **IS effect**<br>**(Fig 8)** | **IS+Rapa**<br>**effect (Fig 8)** | **Expected protein**<br>**effect in normal**<br>**mitophagy\*** |
| --- | --- | --- | --- | --- | --- |
| **PARKIN** | decreased | increased | decreased | reversed | increased |
| **SQSTM1** | decreased | no change | increased | reversed | decreased |
| **ATG7** | no change | no change<br>/decreased | not known | not known | increased |
| **LC3-II** | decreased | not known | decreased | reversed | increased |
| **TOM20** | no change | increased | not known | not known | decreased |

\*. References <sup>34, 60, 76, 77</sup>

## ACKNOWLEDGMENTS

We wish to thank Elaine Seawright, Mahéva Vallet, Bob Fleming, and Graeme Robertson for technical assistance; Rongling Wang, Oliver Lin, and Thomas Tan for experimental help and Yu-Juei Hsu and Lance Chih-Jen Lin for helpful discussions on the manuscript drafts. We thank the staff of the Biological Research Facility (BRF) at the University of Edinburgh for providing animal support, Colin Wood for help with serum analysis and Scott Maxwell for histological assistance. The human femur tissues were collected with the help of Fiona Stewart (research nurse) at the Royal Infirmary of Edinburgh, University of Edinburgh.

## AUTHOR CONTRIBUTIONS

Conceptualisation: SNH, KP, LAS, VEM, KAS and CF. Methodology: SNH, KE, SD, RC and IL. Formal analysis: SNH, CF, KP, TCF, LAS, NMM. Resources: CF, AKA, NMM, TCF. Writing—original draft: SNH, KP and CF. Writing—review and editing: all authors. Visualisation: SNH. Supervision: KAS, LAS, VEM and CF. Funding acquisition: SNH and CF. All authors read and approved the final manuscript.

## SUPPLEMENTARY MATERIAL

### Supplementary Methods

Detailed experimental materials and methods. Unless otherwise stated, all reagents were obtained from Sigma-Aldrich (Dorset, UK).

### Mice

All mice were acclimatised to the conditions of the animal facility until 8 weeks of age before the induction of CKD-MBD. The mice were maintained under standard housing conditions with a 12-hour light-dark cycle and had free access to water and their assigned diet. Body weights (BW) were recorded twice weekly to monitor health conditions until the third week of the study and then daily till sacrifice. Mice losing more than 30% of their BW compared to mice of the control group or reaching the threshold listed in the scoring sheet (Table S1) were euthanised and removed from the study. All animal experiments were approved by the Roslin Institute’s named veterinary surgeon and named animal care and welfare officer (NACWO), with animals maintained in accordance with the UK Home Office code of practice (for the housing and care of animals bred, supplied, or used for scientific purposes). Animal studies were conducted in line with the ARRIVE guidelines. Genotyping of the *mito-*QC mice was performed by diagnostic end-point PCR using genomic DNA isolated from ear notches with the following sets of forward and reverse primers: set 1, 5’-CAA AGA CCC CAA CGA GAA GC-3’ and 5’-CCC AAG GCA CAC AAA AAA CC-3’; and set 2, 5’-CTC TTC CCT CGT GAT CTG CAA CTCC-3’and 5’-CAT GTC TTT AAT CTA CCT CGA TGG-3’ as previously published.^16^

CKD-MBD was induced by feeding a casein-based diet containing 0.6% calcium, 0.9% phosphate and 0.2% adenine (Envigo, Teklad Co. Ltd, Madison, USA). Control (CTL) mice received the same casein-based diet without adenine.^17^

### Human femur bone samples

Approvals were obtained from an ethical committee (Lothian NRS BioResource RTB approval; REC ref-20/ES/0061) in collaboration with the Department of Trauma and Orthopaedic Surgery, The Royal Infirmary of Edinburgh, University of Edinburgh. Patients were recruited before femoral head collection, and samples were collected from patients going through hip fracture surgery (CKD (n = 3) and non-CKD (n = 3) who had given informed consent in writing to the Lothian NRS BioResource. The background information on the human bone samples is listed in Table S2. After removing soft tissue and blood, each bone sample was immersed and washed in phosphate buffered saline (PBS) and stored at -20 C.

### mRNA library preparation

Per-gene read counts were summarised using featureCounts version 1.5.2. Reads were mapped to the mouse genome using the Mus musculus (GRCm38) genome from Ensembl. The annotation used for counting was the standard GTF-format annotation for that reference (annotation version 84). Reads were aligned to the reference genome using STAR (version 2.5.2b),^78^ specifying paired-end reads. Genes were retained in the analysis if they achieved at least 0.1 read counts per million (CPM) in four samples. Reads were normalised using the weighted trimmed mean of the M-values method,^79^ passing ‘TMM’ as the method to the calcNormFactors method of edgeR. Genes were differentially expressed based on fold changes with a 5% false discovery rate (FDR) estimated per the default setting in edgeR between CKD-MBD vs. CTL samples. A fold change cut-off (CKD-MBD vs. CTL, > 1.5-fold) was applied to determine CKD-induced changes.

### Network co-expression and functional enrichment analysis of RNA-seq data

Lists of differentially expressed genes were imported, and a Pearson correlation matrix was constructed using the network analysis program Graphia Professional (https://graphia.app/).^80^ A network graph was created using a threshold value based on a user-defined correlation threshold, where nodes represent transcripts, and the edges represent correlations between transcripts above the threshold. The graph was then clustered into co-expression clusters using the Markov clustering algorithm (MCL) using an inflation value of 1.6, which determined the granularity of clustering.^81^ The differential expressed genes from D20 and D35 contained several genes with different biological functions. We therefore used Fisher’s exact test to determine the probability of a cluster’s composition occurring purely by chance and to statistically confirm the enrichment of a particular cluster related to CKD-MBD. The functional associations of gene clusters were examined through Gene Ontology (GO) enrichment analysis using ToppGene (https://toppgene.cchmc.org/).^82^ Bar plots with histograms of mean expression profiles, each bar representing data from a single mouse bone. GO terms or pathways with a *p* < 0.05 were significantly enriched.

### Serum biochemistry

Serum was collected by centrifugation of the blood at 2500 x g for 10 min at 4°C. Serum concentrations of blood urea nitrogen (BUN), creatinine (Cr), calcium (Ca), and phosphorus (Pi) were quantified immediately using a chemical analyser (Beckman Coulter AU480, Olympus). Enzyme-linked immunosorbent assay (ELISA) kits were used to measure serum concentrations of intact parathyroid hormone (PTH, Pathway Diagnostics, Dorking, UK) and FGF23 (Kainos Laboratories, Inc. Japan). The kits were used according to the manufacturer’s instructions.

### Micro-computed tomography (**μ**CT)

The bone structure of the tibiae was determined using micro-computed tomography (μCT, Skyscan 1172, Bruker, Kontich, Belgium). High-resolution scans with an isotropic voxel size of 5 μm were acquired (60 kV, 167 μA, and 0.5 mm filter, 0.6° rotation angle) and from the reconstructed images (NRecon 1.7.3.0 program; Bruker). The selected trabecular (Tb) volume of interest in the proximal tibial metaphysis began from the bottom of the growth plate, excluding the cortical shell, and extended distally 5% of the entire tibial length. A total of 250 slices beneath this 5% were selected to exclude the primary spongiosa. The cortical bone analysis was performed on datasets derived from μCT scan images at 50% of the total tibial length from the top of the tibia. Bone mineral density (BMD) was calculated as previously described.^17^

### Detection of altered mitophagy in kidneys and bones of *mito*-QC mice

The kidney and femur were fixed in 3.7% paraformaldehyde (PFA), 200 mM HEPES (4-(2-hydroxyethyl)-1-piperazineethanesulfonic acid), pH 7.0. Kidneys were processed to paraffin wax using a Leica ASP300S tissue processor (Leica Microsystems, Milton Keynes, UK). All processed kidneys were sectioned at a 5 μm thickness. Nuclei were counterstained with Hoechst dye (Thermo Fisher Scientific). Images were obtained using an LSM 710 inverted laser scanning confocal microscope (LSCM). Mitolysosome red punta was counted (*mito*-QC Counter) using image analysis software (ImageJ, WI, USA).

### Phalloidin staining of osteocytes from *mito*-QC bone cryosections

The *mito*-QC bone was embedded in optimal cutting temperature compound and cryo-sectioned sagitally. The *mito*-QC tibial bone cryo-section slides were washed in PBS before incubation in 0.1% v/v Triton X-100 in PBS for 20 min at room temperature (RT). The sections were incubated in a solution containing Alexa Fluor 647 Phalloidin (1:500, Thermo Fisher Scientific, Paisley, UK) in the dark for one hour at RT. The slides were then washed, mounted and imaged by a Zeiss LSM 710 inverted LSCM.

### Western blot analysis

The distal and proximal epiphyses of mouse femurs were excised, and the diaphyseal bone marrow was removed by centrifugation at 13,000 x g for 10 min at 4°C. The resultant cortical shafts, human cortical bone samples and cultured osteoblasts were homogenised using a Rotor-Stator Homogenizer (Ultra-Turrax T10) in radioimmunoprecipitation assay (RIPA; Thermo Fisher Scientific) buffer containing protease inhibitors (Roche, Welwyn Garden City, UK). Protein concentrations were determined using the bicinchoninic acid (BCA) protein assay kit (Thermo Fischer Scientific). Equal amounts of extracted proteins were separated using a 4-12% Bis-Tris protein gel (Thermo Fisher Scientific) and then transferred to a nitrocellulose membrane. After blocking in 5% skimmed milk/Tris-buffered saline with Tween 20 (TBST) buffer at RT for 1 hour, the membranes were incubated sequentially with primary and secondary antibodies (Tables S3 and S4). Finally, the protein signals were revealed using the ultra-sensitive enhanced chemiluminescence (ECL) substrates (Thermo Fisher Scientific) and imaged by the GeneGnome XRQ chemiluminescence imaging system (Syngene, UK). Densitometry of the protein bands was analysed with Image J software (NIH) for quantification.

### Cell cultures and viability assay

Calvarial osteoblasts were isolated by sequential enzyme digestion [1 mg/ml collagenase type II (Worthington Biochemical (Lorne laboratories, UK)) in Hanks’ balanced salt solution (HBSS; Life Technologies); 4 mM ethylenediaminetetraacetic acid (EDTA)] and expanded following standard procedures.^17, 83^ The cells were grown in α-minimum essential medium (αMEM, Invitrogen, Paisley, UK) supplemented with 10% fetal bovine serum (FBS) and 0.5% gentamycin (Invitrogen). Upon confluence, mineralisation media supplemented with 50 μg/ml ascorbic acid and 1.5 mM CaCl_2_ were used to grow cells. Where indicated, a range of IS concentrations (0 to 2 mM resuspended in DMSO and BSA) and 75 nM rapamycin was added to the culture medium. Primary cultures were maintained in a 5% CO_2_ atmosphere at 37°C, and the medium was changed every second/third day for up to 7 days. A lactate dehydrogenase (LDH) CytoTox 96 cytotoxicity assay (Promega, Southampton, UK) was performed to assess the dose-dependent effects of IS on cell viability. Cells were treated for up to 14 days, and absorbance intensity was read at 490 nm.

### Mitochondrial staining and membrane potential (ΔΨ) in primary osteoblasts

Primary murine osteoblasts were incubated for 30 min with 100 nM Mitotracker Red CMXRos (Thermo Fisher Scientific), fixed for 15 min in ice-cold methanol, washed, and mounted on glass slides. Cells were examined by a Zeiss LSM 710 LSCM. Quantification of MitoTracker Red staining was assessed at 579□nm.

### Measurement of reactive oxygen species (ROS) production

Mitochondrial ROS production was detected by washing murine primary osteoblasts with a cell-based assay buffer before the mitochondrial ROS detection reagent (Cayman Chemical, Ann Arbor, USA) was added and incubated for 30 mins. The fluorescence was measured using a fluorescence plate reader (Spectromax GEMINI XS, Molecular Devices, Sunnyvale, USA) at excitation 490 nm and emission 560 nm. Cellular ROS was detected by adding 20 μM dihydrofluorescein diacetate (Abcam, Cambridge, UK) to the cells and incubated at RT in the dark for 45 mins. The fluorescence was measured at excitation 485 nm and emission 535 nm.

### Measurement of respiration and glycolysis in osteoblasts challenged by indoxyl sulfate

Primary murine osteoblasts were plated on XF24 microplates (Agilent Technologies) at 1.0×10^4^ cells per well. Upon confluence, cells were treated with and without IS in the medium for 7 days, with changes every two or three days. On the day of respiratory analysis, cells were washed 3 times with Seahorse XF assay medium (Agilent Technologies) supplemented with 25 mM glucose and 10 mM pyruvate. The pH was adjusted with NaOH to 7.35 at 37°C. Cells were then incubated in a CO_2_-free incubator at 37°C for 1 hour and then placed in a Seahorse XFe24 bioanalyser (Agilent Technologies). After basal measurements of oxygen consumption rate (OCR), respiratory ATP synthesis was inhibited by injecting oligomycin (an ATP synthase inhibitor) at a final concentration of 1 μM, causing a drop in OCR representative of ATP synthesis-linked respiration. Next, the electron transport chain (ETC) was uncoupled from ATP synthesis by injecting 0.5, 1.5, and 3 μM final concentration of carbonyl cyanide 4-(trifluoromethoxy) phenylhydrazone (FCCP), increasing OCR to a state of maximal respiration. Finally, respiration was abolished by injection of rotenone and antimycin A (inhibitors of mitochondrial complexes I and III, respectively) at a concentration of 1 μM each. ATP-linked respiratory OCR was calculated from the difference between basal and oligomycin-inhibited OCR. The difference between basal OCR and FCCP-induced OCR was used to calculate the respiratory reserve capacity. The difference in OCR between FCCP and rotenone/antimycin A was considered maximal respiratory capacity. Proton leak respiratory OCR was calculated from the difference between oligomycin-inhibited OCR and rotenone/antimycin A-inhibited OCR. The increase in ECAR after the addition of glucose is representative of the cellular glycolytic flux. Oligomycin was injected to a final concentration of 1 μM, inhibiting respiration-derived ATP, leading to an increase in ECAR. This increase was considered the glycolytic reserve. Maximal glycolysis was also estimated. All data were normalised to total protein concentration by sulforhodamine B staining.

### Immunofluorescence of primary osteoblasts

Immunofluorescence (IF) localisation of mitophagy-associated proteins was performed on primary osteoblasts cultured on sterilised glass coverslips in the 24-well plate at a density of 2.5 x 104 cells/cm^2^. Cells were treated with IS (2 mM) and/or rapamycin (75 nM) for 7 days. To accurately evaluate autophagy, autophagic flux (puncta) was measured via the inhibition of lysosomal degradation by bafilomycin (Baf, 10 nM) treatment for 6 hrs prior to the end of the experiment. Cells were then fixed in 4% PFA for 20 min, permeabilised for 10 min, and blocked in PBS containing 5% goat serum at RT for 1 h. The cells were incubated overnight at 4□ with primary antibodies (Table S2) in PBS buffer containing 5% normal goat serum. After washing with PBS, AlexaFluor-conjugated secondary antibodies (Table S5) made up in PBS were applied for 1 hr at RT. Nuclei were counterstained with Hoechst. Cells were mounted and imaged by LSM 710 LSCM.

## SUPPLEMENTARY FIGURE LEGENDS

**Supplementary Figure S1. Adenine-induced CKD-MBD mice are osteopenic and show altered biomechanical properties of the tibia.** (a) CKD-MBD mice exhibited decreased trabecular (Tb) bone mineral density (BMD), separation (Sp), connectivity density (Conn Dn), structural model index (SMI), and pattern factor (Pf) at D35. (b) CKD-MBD mice had increased cortical (Ct) BMD and closed porosity (Po (cl)) but decreased bone volume/tissue volume (Ct BV/TV), thickness (Ct Th), and cross-sectional area (Ct CSA) at a region 50% of the total tibial length. (c) Representative 3D micro-CT images of trabecular and cortical bone from CTL and CKD-MBD mice at D35. (d) Biomechanical properties of the tibia from CTL and CKD-MBD mice indicated that maximum load, load at rupture, and stiffness were reduced in the CKD-MBD bones. Tibia of n ≥ 3 biological replicates were analysed. The data were represented as the means ± SEM. NS, not significant, * *p* < 0.05; ** *p* < 0.01 vs. CTL mice of the same age.

**Supplementary Figure S2. Representative bar plots with histograms of mean expression profiles from the *Fgf23* cluster.** Each bar represents data from a single mouse femur (4 bars of each time point, from left to right, CTL, yellow: D0, D20, D35, and then CKD-MBD, green: D5, D10, D15, D20, D25, D30, D35). All genes (*Fgf23*, *Cma1*, *Mcpt4*, *Cxcl13*, *Tpsb2*) within the same cluster (Cluster 16) had similar expression patterns by time and biological function.

**Supplementary Figure S3. Osteocytes of CKD-MBD mice have a disorganised dendritic network.** (a) Representative confocal images of the osteocyte dendritic network from the cortical bone of CTL and CKD-MBD *mito*-QC mouse femurs. The interconnected phalloidin-labelled dendrites were reduced in CKD-MBD osteocytes. (b) The total surface area of phalloidin-stained dendrites was analysed. Each dot represents the dendrites’ total surface area (μm^2^)/three osteocytes/section. Data were presented as means ± SEM (4 sections/mouse, n = 3). Scale bar 10 μm. Analysis was tested by a Student’s t-test. **** *p* < 0.0001 vs the CTL group.

**Supplementary Figure S4. Dietary adenine treatment induced mitophagy in the kidneys of *mito*-QC mice.** (a) Kidneys from *mito*-QC mice fed a CTL or adenine-supplemented diet were imaged at D35. Representative confocal images of the renal cortex and glomerulus are shown. Scale bar 50 μm (renal cortex) and 10 μm (glomerulus). (b) Higher magnifications show representative images of red puncta (white arrows) in the glomerulus, proximal collecting tubule (PCT), and distal collecting tubule (DCT). Scale bar 10 μm. (c) Quantification of red mitophagy puncta within the glomerulus, PCT and DCT. Two tissue sections from each *mito*-QC mouse (n = 3) and a total of 9 glomeruli, 12 PCTs, and DCTs were analysed. Data are presented as mean ± SEM. Analysis was done by Student’s t-test. * *p* < 0.05; **** *p* < 0.0001; vs. respective CTL.

**Supplementary Figure S5. Assessment of toxic effects of IS on primary osteoblasts.** (a) Cells were exposed to various concentrations of indoxyl sulfate (IS; 0, 0.25, 0.5, 1, and 2 mM) for 7 days, (b) 8 days, (c) 10 days, (d) 11 days, (e) 12 days, and (f) 14 days. Cell cytotoxicity was increased after 12 days of 2 mM IS treatment. The data are represented as the mean ± SEM (n = 4), * *p* < 0.05, vs. non-treated CTL group (0 mM).

**Supplementary Figure S6. Localisation of PARKIN expression in primary osteoblasts treated with IS and rapamycin.** Primary osteoblasts were treated with 0 or 2 mM indoxyl sulfate (IS; 2 mM) ± rapamycin (rapa;75 nM) for 7 days. Red: PARKIN; green: TOM20; blue: Hoechst, Scale bar, 10 μm.

**Supplementary Figure S7. Localisation of p62/SQSTM1 expression in primary osteoblasts.** Primary osteoblasts were treated with 0 or 2 mM indoxyl sulfate (IS; 2 mM) ± rapamycin (rapa; 75 nM) for 7 days. Red: SQSTM1; green: TOM20; blue: Hoechst, Scale bar, 10 μm.

## SUPPLEMENTARY TABLES

**Supplementary Table S1.** Animal health assessment sheet for CKD-MBD model.

|  |  |  |  |  |  |  |  |  |
| --- | --- | --- | --- | --- | --- | --- | --- | --- |
| Mouse ID: |  |  |  |  |  |  |  | PPL # |
| **Date:** |  |  |  |  |  |  |  |  |
| **APPEARANCE** |  |  |  |  |  |  |  |  |
| Moderate piloerect |  |  |  |  |  |  |  |  |
| Poor body condition, *e.g.*,<br>square tail |  |  |  |  |  |  |  |  |
| Failure to groom |  |  |  |  |  |  |  |  |
| Hunched posture |  |  |  |  |  |  |  |  |
| **BEHAVIOR** |  |  |  |  |  |  |  |  |
| Isolated, away from others |  |  |  |  |  |  |  |  |
| Restless |  |  |  |  |  |  |  |  |
| Reluctant to move |  |  |  |  |  |  |  |  |
| Inappetence |  |  |  |  |  |  |  |  |
| Motor behavioural<br>abnormalities |  |  |  |  |  |  |  |  |
| **CLINICAL SIGNS** |  |  |  |  |  |  |  |  |
| Diarrhoea |  |  |  |  |  |  |  |  |
| Breathing abnormal |  |  |  |  |  |  |  |  |
| Initials |  |  |  |  |  |  |  |  |
| Date |  |  |  |  |  |  |  |  |

1. Place the numerical score in the appropriate box (see below)
If there is cause for concern, inform the NACWO, NVS, and/or the PPL holder and record details in the “Animal Room Log Book.”
2. Score System: normal = 0, slightly abnormal = 1, significant change = 2
3. Euthanasia will be performed when the following criteria are met: (1) A score of 2 is given to 4 or more parameters (except for “clinical signs”), (2) A total score of 7 is reached from any parameters, (3) Both clinical signs receive a score of 1, (4) An overall weight loss of 30% is observed.

**Supplementary Table S2.** Background information on the human bone samples.

| No. of samples | Group | Gender | Age | eGFR | Underlying diseases |
| --- | --- | --- | --- | --- | --- |
| **1** | Control | M | 53 | > 60 | Depression, history of gastric ulcer and pneumonia |
| **2** | Control | F | 78 | > 60 | Gastritis |
| **3** | Control | F | 86 | > 60 | History of deep vein thrombosis |
| **4** | CKD | M | 80 | 47 | Parkinson's disease, paroxysmal atrial fibrillation, DVT |
| **5** | CKD | F | 83 | 28 | Hypertension, osteoporosis |
| **6** | CKD | F | 89 | 39 | Atrial fibrillation |

**Supplementary Table S3.**
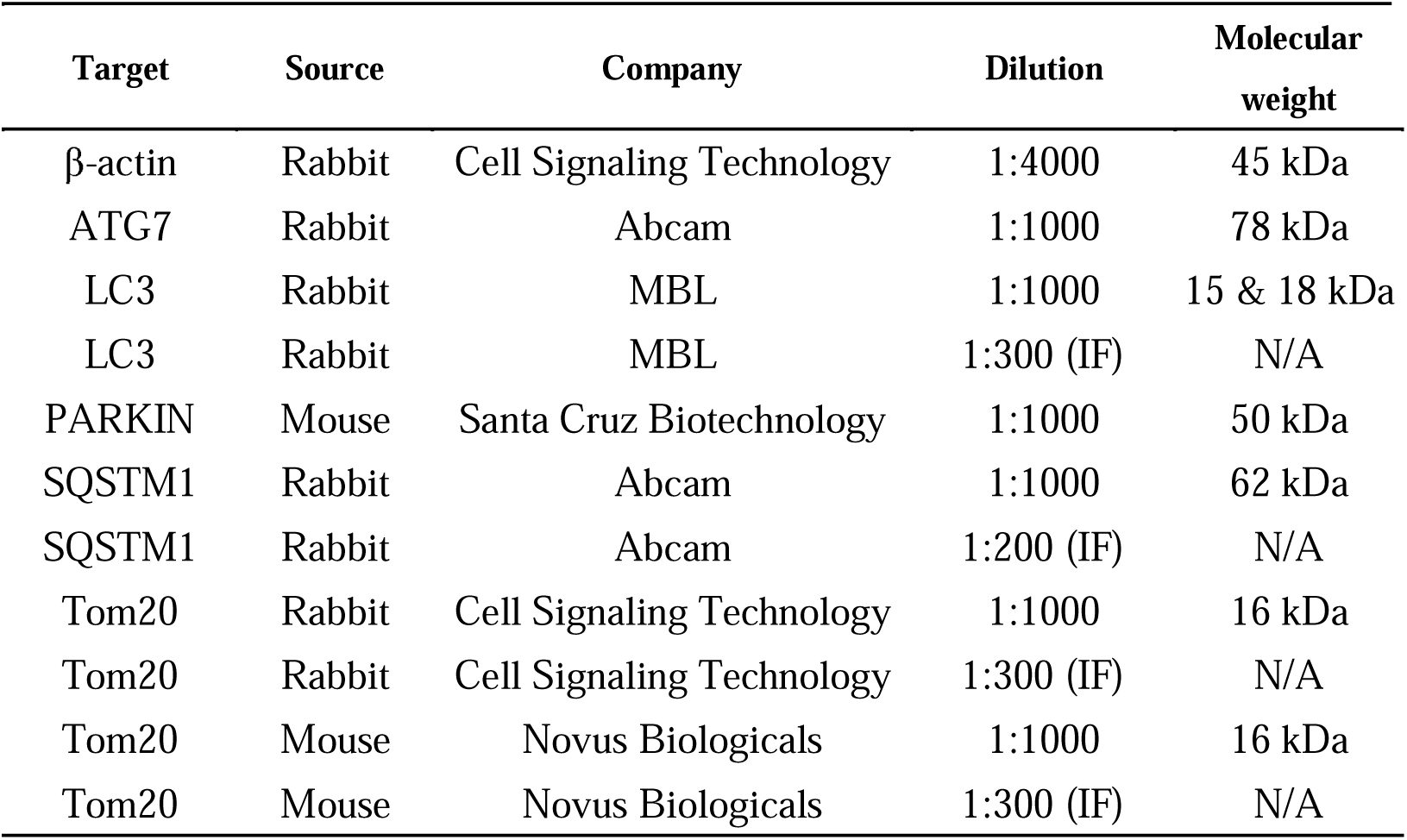
Primary antibodies used for western blotting and immunofluorescence.

**Supplementary Table S4.** Secondary antibodies used for western blotting.

| Target | Source | Company | Dilution |
| --- | --- | --- | --- |
| β-actin | Goat anti-rabbit | Dako | 1:1000 |
| ATG7 | Goat anti-rabbit | Dako | 1:1000 |
| LC3 | Goat anti-rabbit | Dako | 1:1000 |
| PARKIN | Goat anti-mouse | Dako | 1:1000 |
| SQSTM1 | Goat anti-rabbit | Dako | 1:1000 |
| Tom20 | Goat anti-rabbit | Dako | 1:1000 |
| Tom20 | Goat anti-mouse | Dako | 1:1000 |

**Supplementary Table S5.**
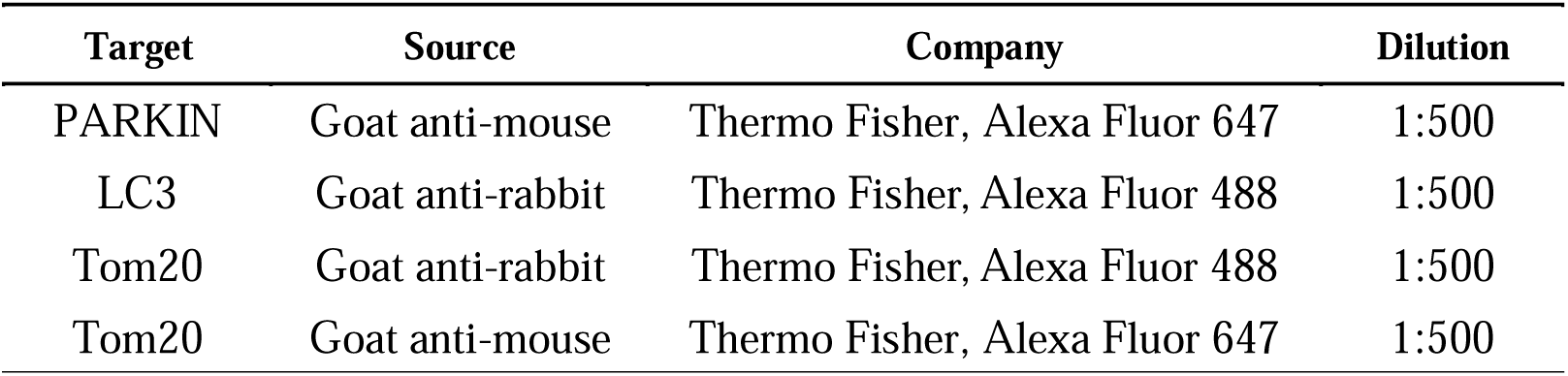
Secondary antibodies used for immunofluorescence.

| Target | Source | Company | Dilution |
| --- | --- | --- | --- |
| PARKIN | Goat anti-mouse | Thermo Fisher, Alexa Fluor 647 | 1:500 |
| LC3 | Goat anti-rabbit | Thermo Fisher, Alexa Fluor 488 | 1:500 |
| Tom20 | Goat anti-rabbit | Thermo Fisher, Alexa Fluor 488 | 1:500 |
| Tom20 | Goat anti-mouse | Thermo Fisher, Alexa Fluor 647 | 1:500 |

## ADDITIONAL INFORMATION

**Supplementary Additional Data File 1:** The differentially expressed genes (1.5-fold up and down, *p* < 0.05) in a comparison between CKD and control-treated mice from time points of D20 and D35.

