## Supplementary figures and images for "Mitochondrial dysfunction and mitophagy blockade contribute to renal osteodystrophy in chronic kidney disease-mineral bone disorder"

### Figure S1a-b

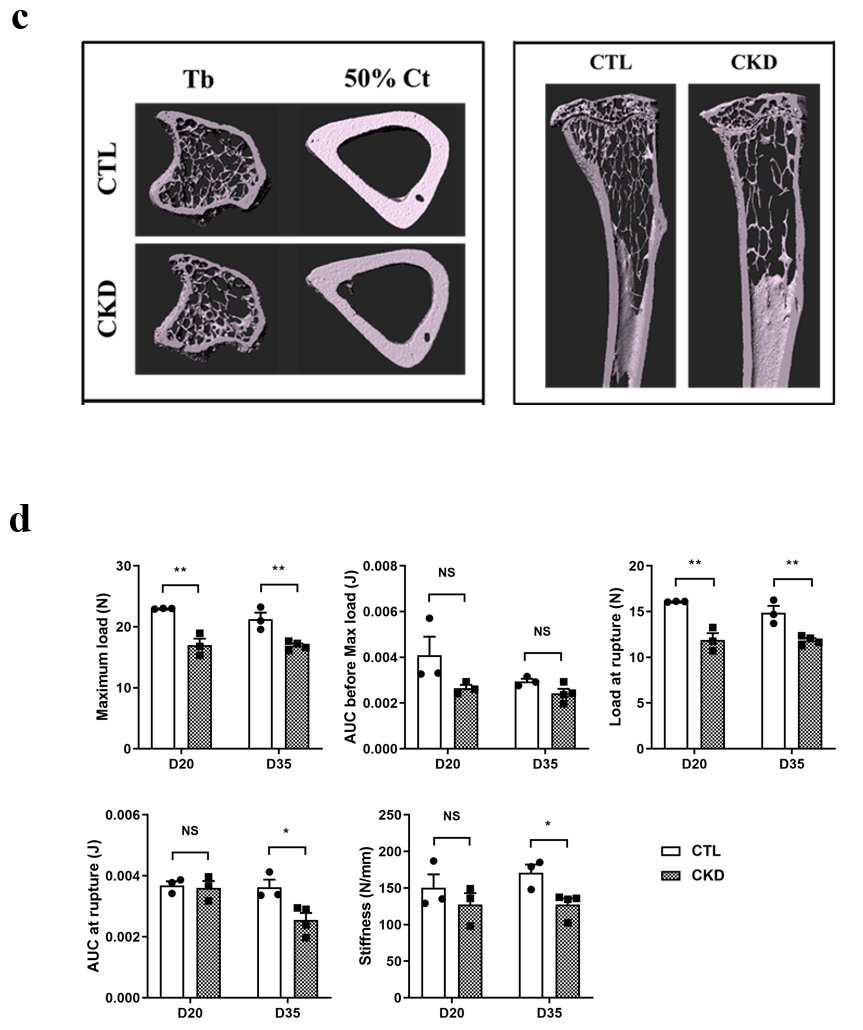

### Figure S1c-d

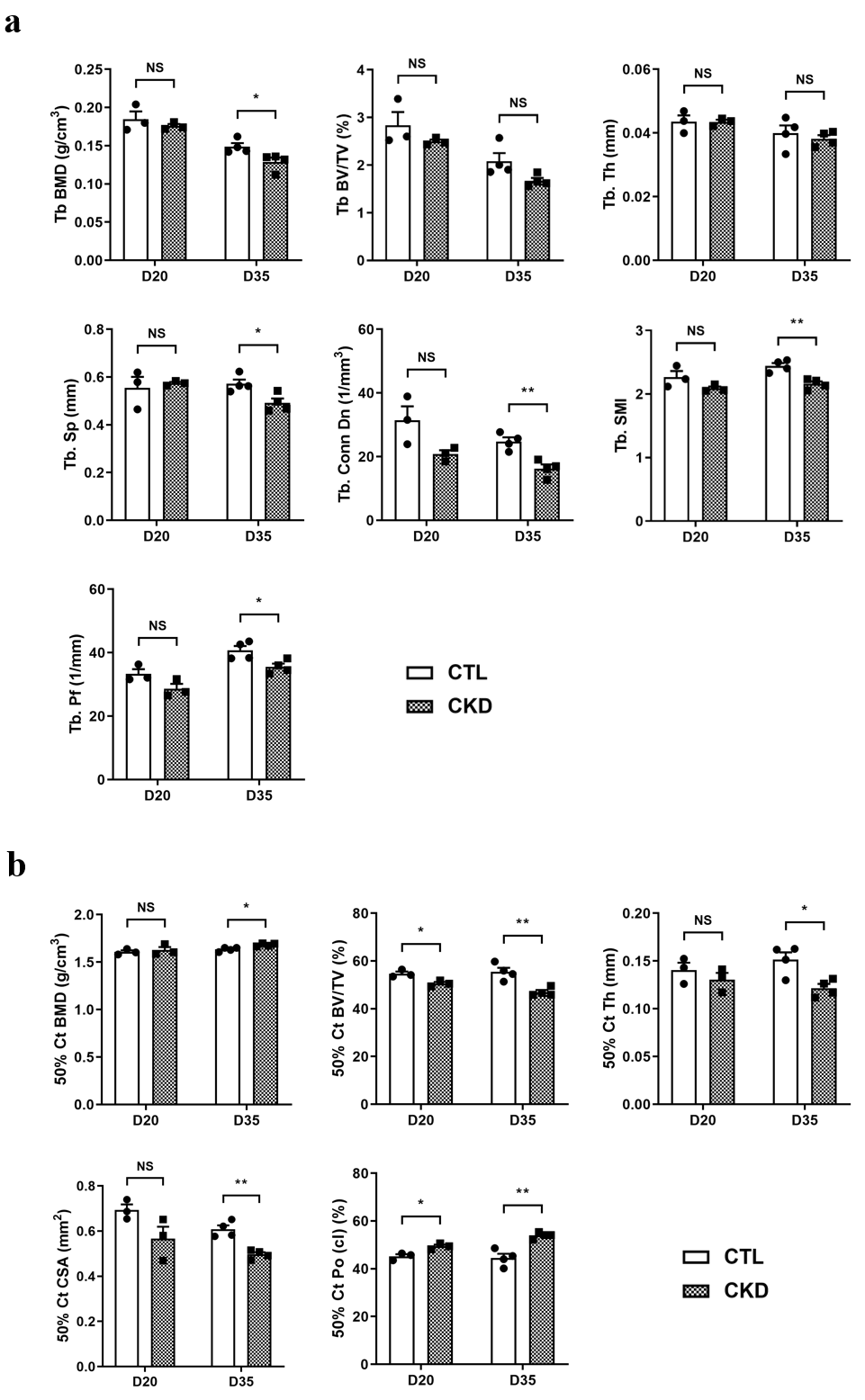

### Figure S2

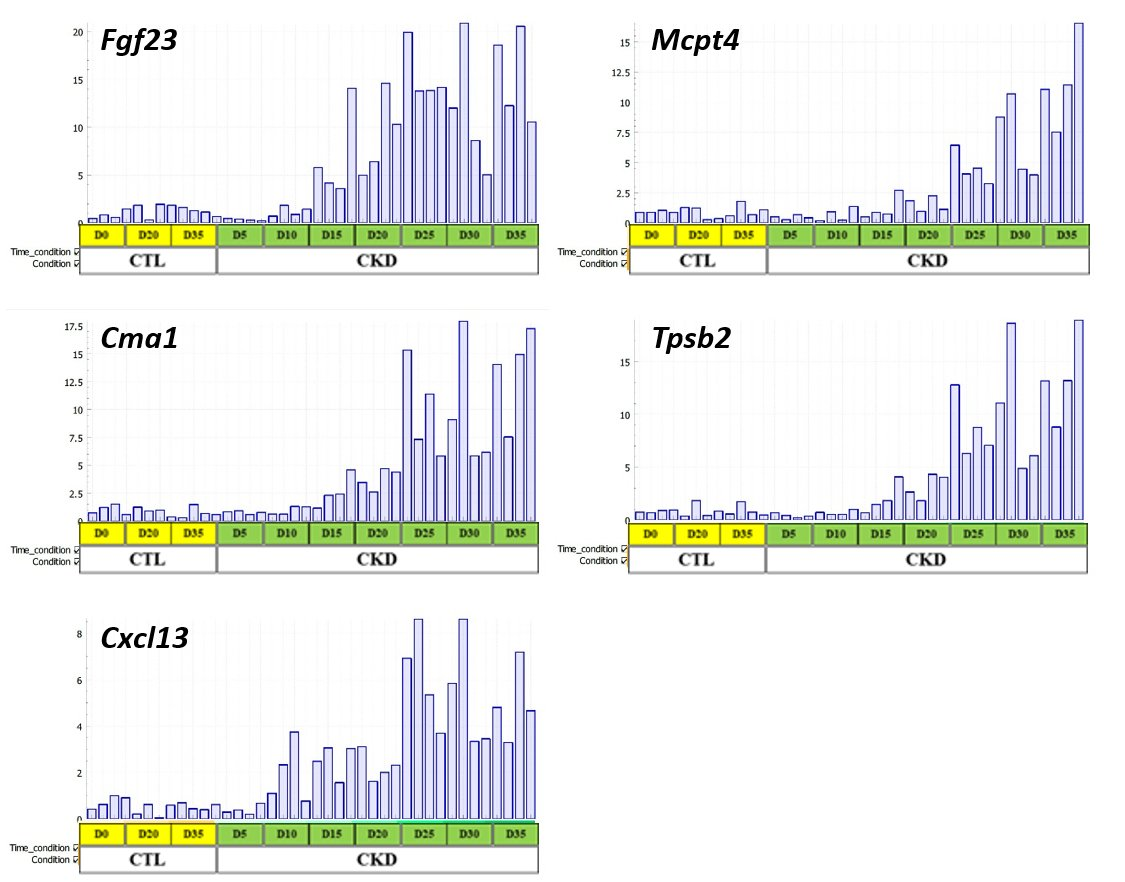

### Figure S3

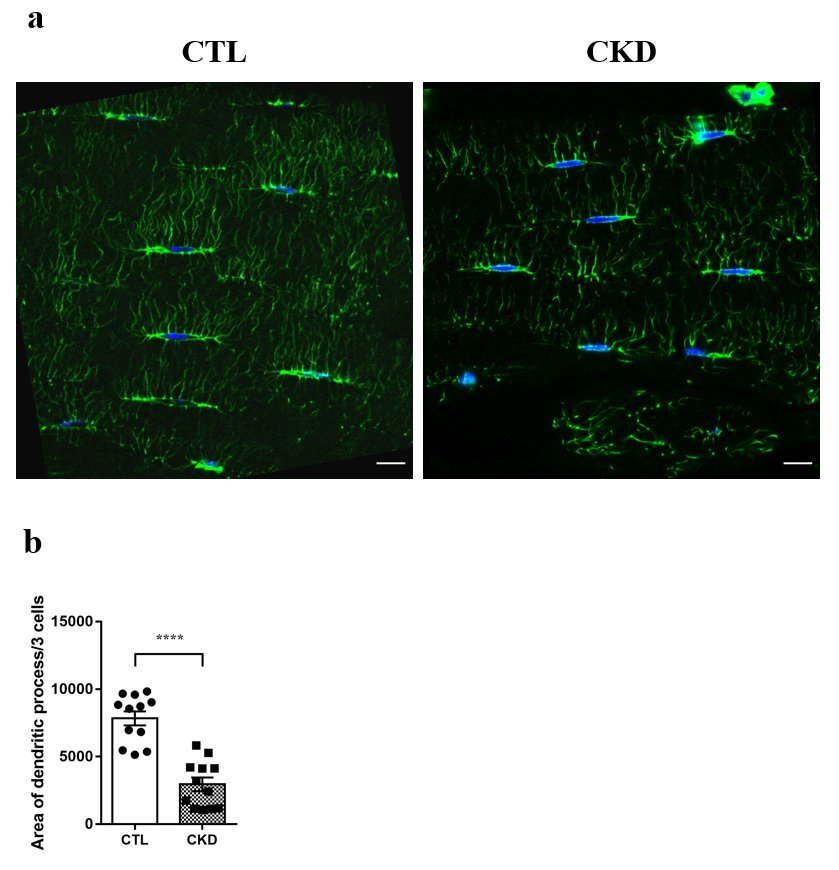

### Figure S4

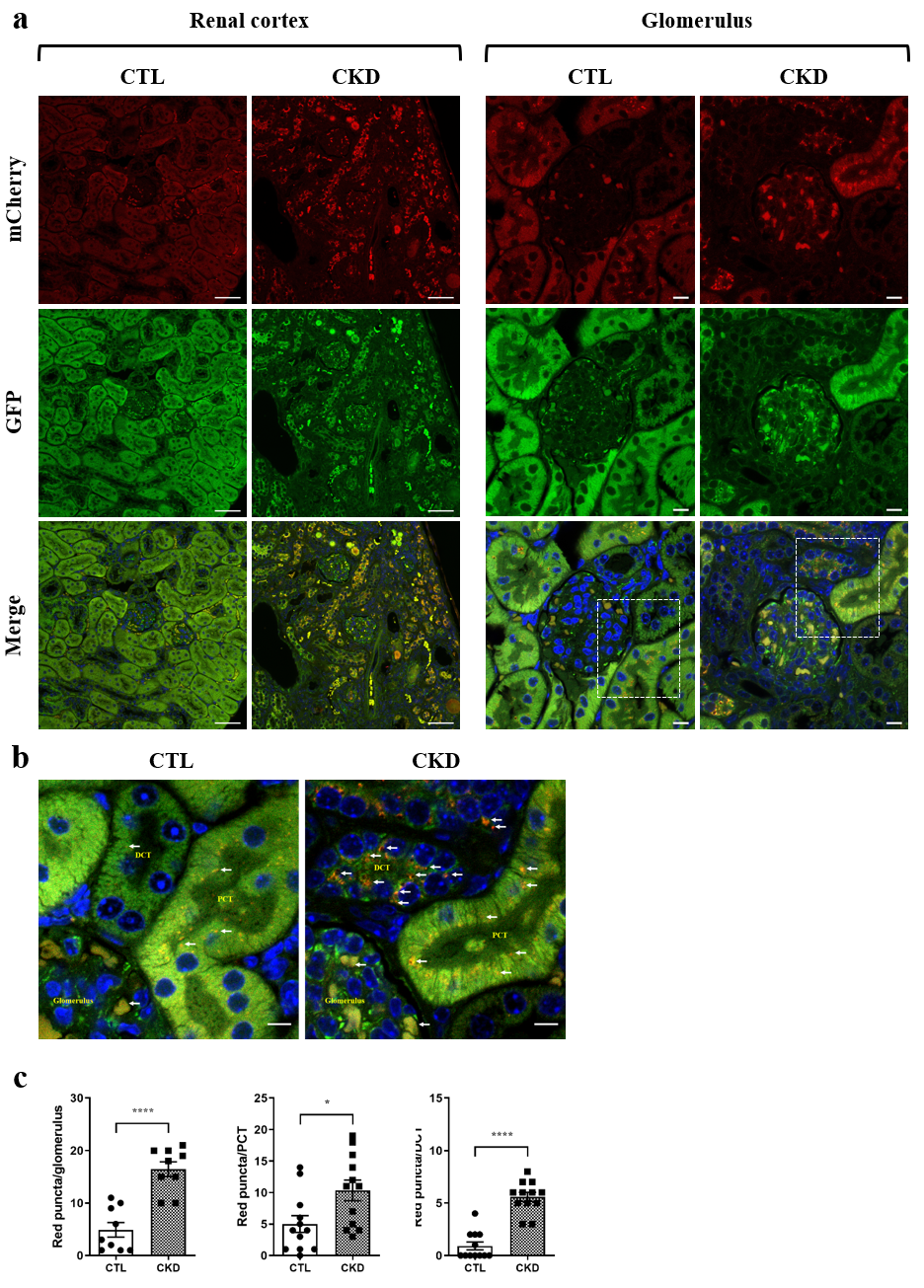

### Figure S5

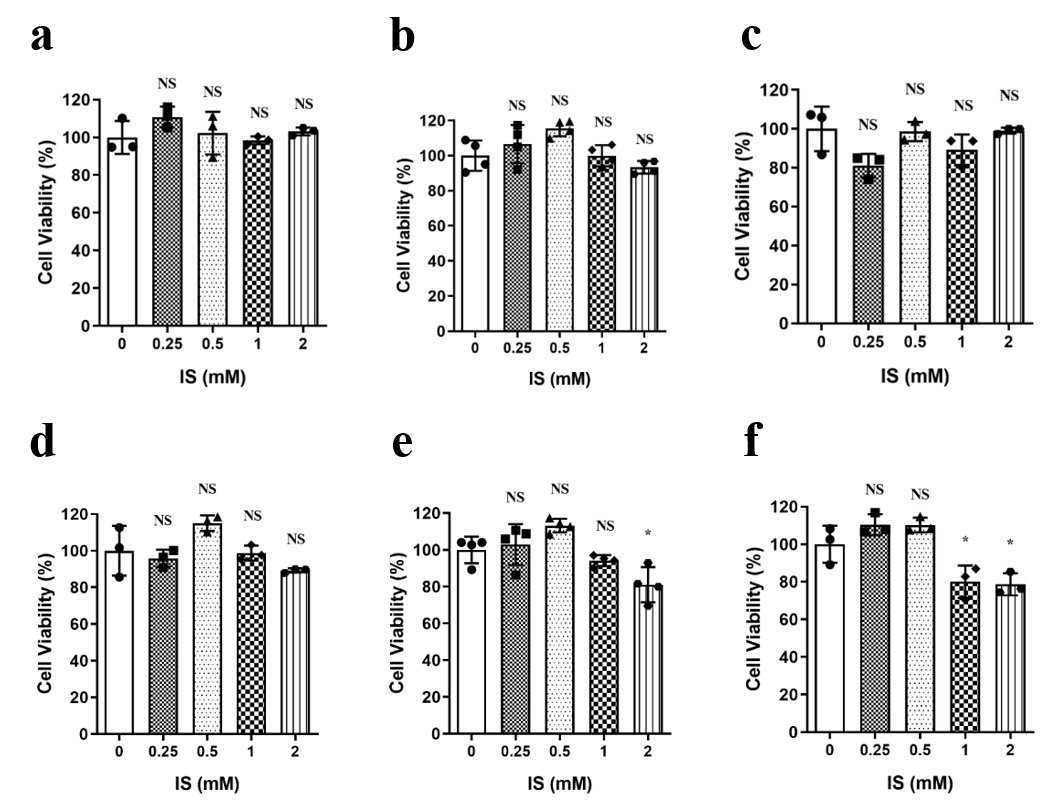

### Figure S6

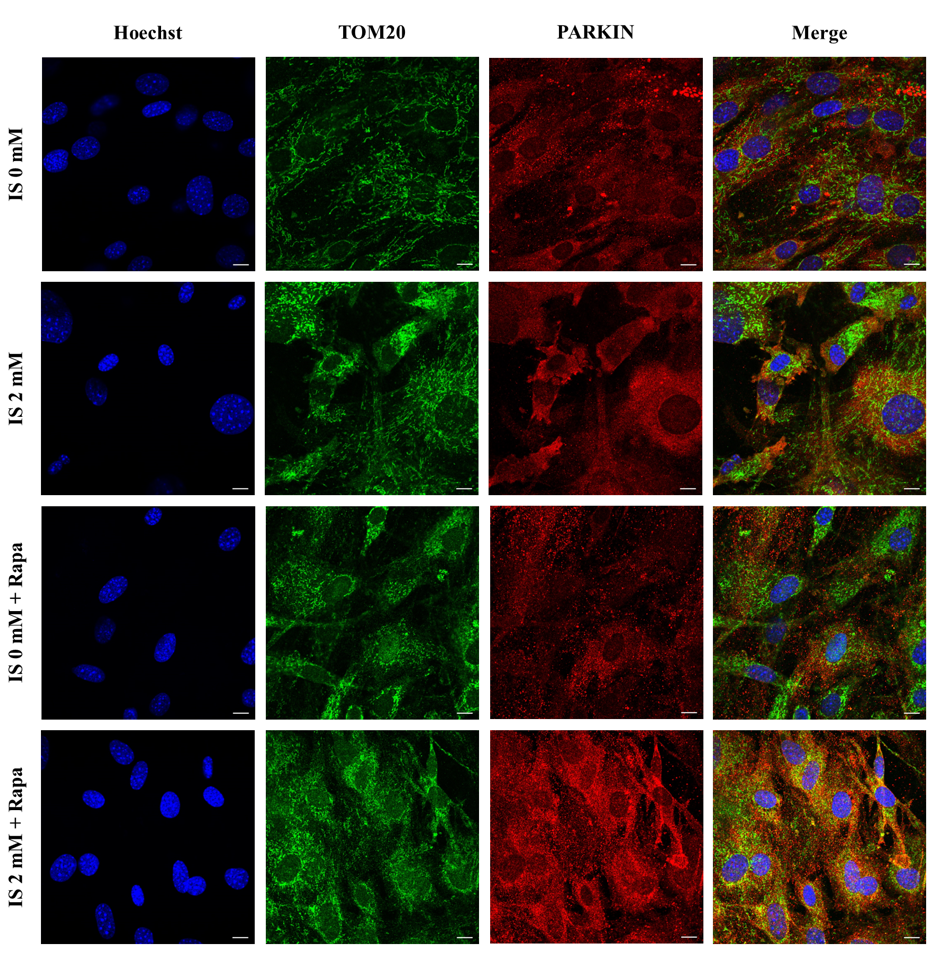

### Figure S7

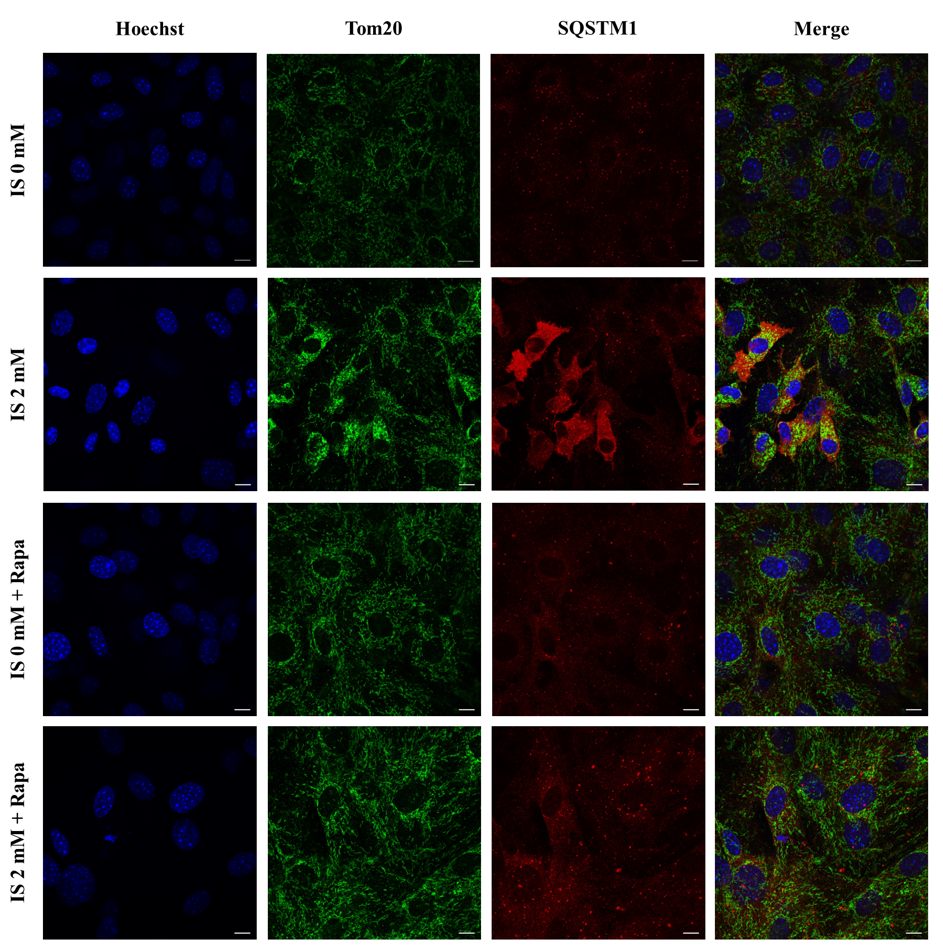
